# Genetic correlates of evolutionary adaptations in cognitive functional brain networks and their relationship to human cognitive functioning and disease

**DOI:** 10.1101/671610

**Authors:** Yongbin Wei, Siemon C. de Lange, Lianne H. Scholtens, Kyoko Watanabe, Dirk Jan Ardesch, Philip R. Jansen, Jeanne E. Savage, Longchuan Li, Todd M. Preuss, James K. Rilling, Danielle Posthuma, Martijn P. van den Heuvel

**Affiliations:** Connectome Group, Department of Complex Trait Genetics, Center for Neurogenomics and Cognitive Neuroscience, Amsterdam Neuroscience, Vrije Universiteit Amsterdam, Amsterdam, The Netherlands; Department of Complex Trait Genetics, Center for Neurogenomics and Cognitive Neuroscience, Amsterdam Neuroscience, Vrije Universiteit Amsterdam, Amsterdam, The Netherlands; Department of Child and Adolescent Psychiatry, Erasmus Medical Center, Rotterdam, The Netherlands; Marcus Autism Center, Children’s Healthcare of Atlanta, Emory University School of Medicine, Atlanta, GA, USA; Division of Neuropharmacology and Neurologic Diseases, Emory University, Atlanta, GA, USA; Center for Translational Social Neuroscience, Emory University, Atlanta, GA, USA; Yerkes National Primate Research Center, Emory University, Atlanta, GA, USA; Department of Anthropology, Emory University, Atlanta, GA, USA; Center for Behavioral Neuroscience, Emory University, Atlanta, GA, USA; Department of Psychiatry and Behavioral Sciences, Emory University, Atlanta, GA, USA; Department of Clinical Genetics, Amsterdam University Medical Center, Amsterdam Neuroscience, Amsterdam, The Netherlands

**Keywords:** functional networks, cortical expansion, gene expression, human brain evolution, GWAS, intelligence, social cognition, Schizophrenia, psychiatric disorder

## Abstract

Cognitive functional networks such as the default-mode network (DMN), frontal-parietal network (FPN), and salience network (SN), are key networks of the human brain. Here, we show that the distinct rapid evolutionary cortical expansion of cognitive networks in the human brain, and most pronounced the DMN, runs parallel with high expression of genes important for human evolution (so-called HAR genes). Comparative gene expression examination then shows that HAR genes are more differentially expressed in cognitive networks in humans compared to the chimpanzee and macaque. Genes with distinct high expression in the DMN display broad involvement in the formation of synapses and dendrites. Next, we performed a genome-wide association analysis on functional MRI data, and show that HAR genes are associated with individual variations in DMN functional connectivity in today’s human population. Finally, gene-set analysis suggests associations of HAR genes with intelligence, social cognition, and mental conditions such as schizophrenia and autism. Taken together, our results indicate that the expansion of higher-order functional networks and their cognitive properties have been an important locus of change in recent human brain evolution.

## Introduction

The human brain is widely believed to be able to support more complex cognitive abilities than other highly developed and intelligent primate species, such as the chimpanzee, with which we share the majority of our genetic material^1^. This discrimination in cognitive abilities is thought to be related to the rapid expansion of multimodal association areas and their structural and functional connections in the human brain^2, 3, 4^, with cognitive functional networks, such as the frontal-parietal network (FPN), salience network, and default-mode network (DMN), to play an essential role in higher-order cognitive brain functions^5, 6, 7^. These cognitive functional networks are known to be highly heritable^8^ and to relate to genetic effects associated with neuron growth and metabolic effects^9^, suggesting considerable genetic control of these higher-order cognitive systems. Understanding the evolutionary genetic underpinnings of these cognitive functional networks, and in particular, to what extent these functional networks have been developed in recent human evolution, is crucial for our understanding of the enhancement of complex brain functions in humans.

The DMN specifically includes a spatially distributed functional network of posterior cingulate, precuneus, inferior parietal, middle temporal, and medial prefrontal cortices^5, 6^. Comparative neuroimaging studies have shown default-mode activity in chimpanzees^10^ and macaques^11^, but with potential subtle differences in both the spatial and topological layout of this central functional network in comparison to humans^12^, changes that may relate to enhancement of cognitive functions in humans compared to other primate species. The DMN is central to social cognition, including aspects of mental self-projection^13^ to rehearse future actions mentally^14^ and to understand another person’s mental perspective (theory of mind^15^). These advanced social abilities are believed to be more developed from those of other primate species and were likely highly adaptive during recent human evolution^16^, enabling humans to potentially make more complex social inferences and to more accurately anticipate beliefs and actions of others^13^.

Here, we set out to study the role of evolutionary genetic control of higher-order cognitive networks in recent human brain evolution, with a particular focus on the evolutionary changes of the DMN. Recent genome-wide studies comparing the human genome with that of the chimpanzee have identified a unique set of loci that displayed accelerated divergence in the human lineage^17, 18^. Genes associated with these so-called human accelerated regions (HAR) have been linked to neuron development, but also the development of brain disorders such as autism spectrum disorder (ASD)^19^. By integrating data from comparative neuroimaging and genetics, we first demonstrate that regions of higher-order cognitive networks are highly expanded in recent human brain evolution. We next show that HAR genes are highly expressed in expanded regions of higher-order cognitive networks, particularly in regions of the DMN, and HAR genes are more differentially expressed in regions of higher cognitive networks in the human brain than in comparable regions in the chimpanzee and macaque brain. By means of a genome-wide association study (GWAS), we then show evidence of a significant role of evolutionary HAR genes in DMN functional connectivity in today’s human population. We show that sets of HAR and DMN genes are significantly involved in human intelligence, social behavior, and mental disorders. Altogether, our findings provide evidence of genes important for human evolution to have played a central role in shaping the functional cognitive brain network architecture of modern-day humans.

## Results

### Human cortical expansion

We started by examining the expansion of the human cortex (*Homo sapiens*) compared to the cortex of the chimpanzee (*Pan troglodytes*), one of our closest living evolutionary relatives along with bonobos (*Pan paniscus*). We particularly hypothesized a large expansion in the human brain of cortical areas involved in higher-order cognitive networks. Cortical morphometry of the chimpanzee and human cortex was assessed by means of a surface-to-surface mapping of 3D reconstructions of the cortical mantle across both species, based on *in vivo* T1-weighted MRI (29 chimpanzees, 30 humans; see Methods; Fig. 1a). The largest expansion of the human cortex was found in areas of bilateral orbital inferior frontal gyrus (×4.0 expansion), rostral middle frontal lobe (×3.8 expansion), inferior/middle temporal lobe (×3.0/2.9 expansion), lingual gyrus (×2.9 expansion), right inferior parietal lobe (×3.7 expansion), and left precuneus (×2.7 expansion; adjusted *p* < 0.001, false discovery rate [FDR] corrected; Cohens’*d* > 0.989; see Fig. 2a and Supplementary Table 1). The lowest expansion was found in primary areas, including bilateral precentral gyrus (×1.3 expansion), postcentral gyrus (×1.4 expansion), and paracentral lobe (×1.2 expansion; adjusted *p* < 0.001, FDR corrected; Cohens’*d* < −1.047; see Fig. 2a and Supplementary Table 1). A relative low expansion was also found in middle/posterior cingulate cortex (×1.5 expansion, adjusted *p* < 0.001), which might be attributable to the involvement of cingulate cortex in the paleomammalian brain that arose early in mammalian evolution^20^.

**Figure 1.**
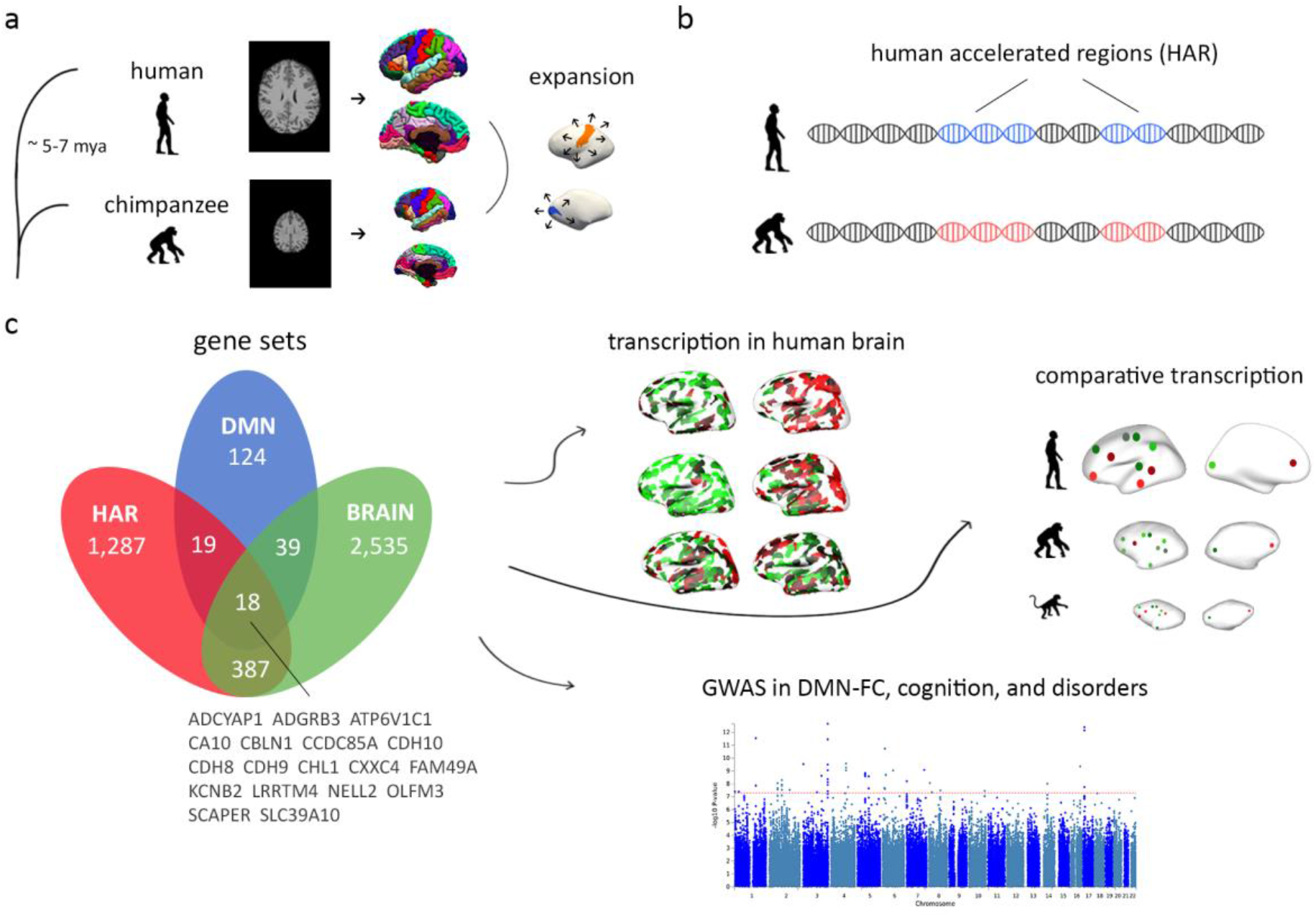
Overview of our methods. (a) The human and chimpanzee cortex are reconstructed using MRI data, and chimpanzee-to-human cortical expansion is computed based on the reconstructed cortex. (b) Genes associated with human accelerated regions (HAR), which represent genomic loci with accelerated divergence in humans, are examined in the current study. (c) Cortical gene expression of HAR genes and HAR-BRAIN genes (as HAR genes that are preferentially expressed in the brain) are examined using human transcription data from the Allen Human Brain Atlas (AHBA) and comparative transcription data of the human, chimpanzee, and macaque from the PsychENCODE database. HAR, HAR-BRAIN, and DMN genes (as a set of genes that preferentially express in the default-mode network) are further linked to genetic variants associated with the DMN functional connectivity, cognitive abilities, and disorders provided by genome-wide association study (GWAS).

**Figure 2.**
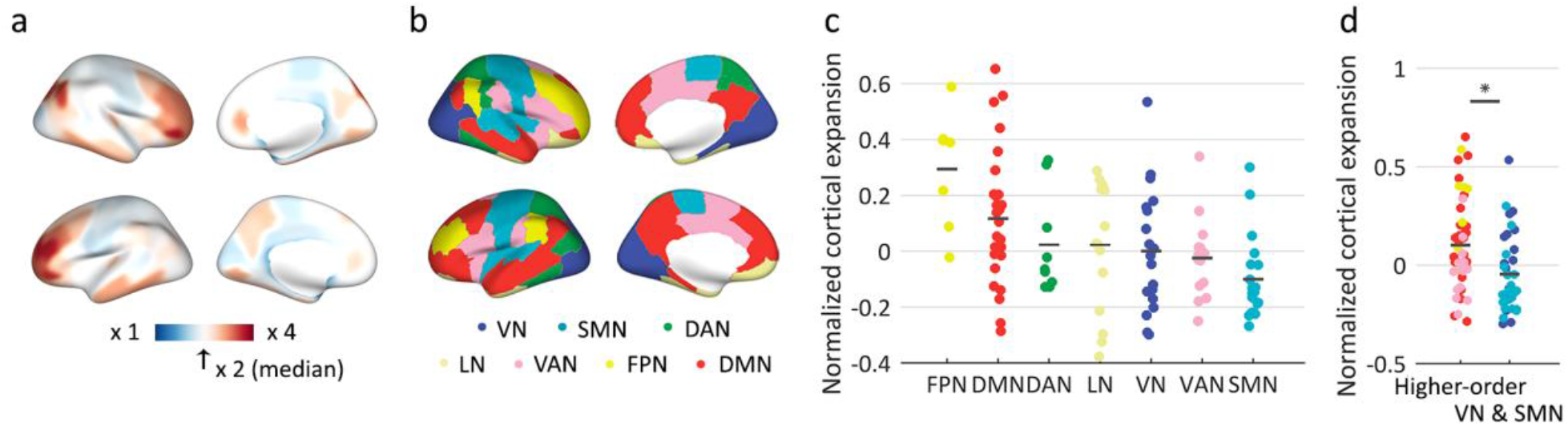
Cortical expansion. (a) Cortical expansion from chimpanzees to humans. Blue: below-median human expansion (i.e., < ×2 compared to chimpanzee); red: above-median expansion (i.e., > ×2). (b) Brain map of the seven resting-state functional networks according to the DK-114 atlas, describing the visual (VN), somatomotor (SMN), dorsal-attention (DAN), limbic (LN), ventral-attention (VAN), frontal-parietal (FPN), and default mode network (DMN). (c) Levels of normalized cortical expansion per functional network in descending order of mean expansions. (d) Levels of normalized cortical expansion in higher-order networks (DMN, FPN, VAN) versus VN and SMN. Dots depict cortical regions. Colors represent functional networks, as in (b). Central marks are mean expansions. * indicates a two-sided *p* < 0.05, FDR corrected.

Since we were particularly interested in examining potential differential expansion of cognitive functional networks compared to primary functional networks, we next used the functional brain atlas of Yeo and colleagues^21^ to assign all cortical areas into the visual (VN), somatomotor (SMN), dorsal-attention (DAN), limbic (LN), ventral-attention (VAN, also commonly referred to as the salience network), frontal-parietal (FPN, also commonly referred to as central-executive network), and default-mode network (DMN) (Fig. 2b). Higher-order cognitive functional networks (i.e., DMN, FPN, VAN) displayed particularly high levels of cortical expansion as compared to the SMN/VN (×1.2 larger expansion in regions of cognitive networks combined compared to the regions of the SMN/VN combined, *t*(86) = 3.257, *p* = 0.002; Fig. 2d). Furthermore, we found regions of the FPN to show the largest expansion (mean: ×2.9 expansion), with the DMN in second place (mean: ×2.4 expansion), showing both significantly higher expansion when comparing each of them with the rest of the brain (FPN: *t*(108) = 3.360, *p* = 0.001; DMN: *t*(108) = 2.621, *p* = 0.010; FDR corrected; Supplementary Table 2). In contrast, examining the other five networks, separately, did not show significant increases in the expansion of these networks compared to the rest of the cortex.

### HAR gene expression

We continued by examining this distinct pattern of human cortical expansion across the seven resting-state functional networks in relationship to cortical gene expression patterns relevant to human evolution. Microarray data on the spatial pattern of gene expression across cortical regions were obtained from the Allen Human Brain Atlas (AHBA) (http://human.brain-map.org/), containing transcriptome expression profiles of 20,734 genes across 57 areas of the left cortical mantle (see Methods; other atlas resolutions showed similar findings, see Supplementary Results). Genes relevant to human evolution were taken as the list of 2,143 genes associated with human accelerated regions (HAR) of the human genome as presented previously by Doan and colleagues^19^, with HAR-associated genes selected based on positional mapping. Alternative selection and allocation of HAR-associated genes are possible (e.g., using chromatin interactions) and we examined such alternatives to validate our results (Supplementary Results).

From the set of 2,143 HAR-associated genes, expression data of 1,711 genes was present in AHBA (from now on referred to as HAR genes; Supplementary Table 3), and we further examined their cortical gene expression levels in comparison to the comparative cortical expansion. The expression profile of HAR genes was positively associated with the global pattern of human cortical expansion (*r*(53) = 0.359, *p* = 0.008; Supplementary Fig. 1), indicating a variable expression of HAR genes across cortical areas, with in particular the highest expression of HAR genes in areas with large expansion of the human cortex. This observed association between cortical HAR gene expression and cortical expansion significantly exceeded the null condition of correlations between cortical expansion and expression of random gene sets (i.e., selecting 1,711 random genes) selected out from a pool of 8,686 genes related to general evolutionary conserved genetic elements (further referred to as ECE genes; *p* < 0.001, 10,000 permutations; Supplementary Fig. 1)^22^. Cortical regions of the cognitive networks showed significantly higher expression of HAR genes compared to regions of the SMN/VN (*t*(44) = 2.742, *p* = 0.009; FDR corrected; Supplementary Fig. 1), with regions of the DMN showing the highest HAR gene expression (*t*(55) = 2.274, *p* = 0.027, not corrected, when comparing the DMN to all other networks combined; Supplementary Table 4). These effects were also found to significantly exceed the null condition of effects of similar sized sets of random ECE genes (*p* < 0.001, 10,000 permutations; Supplementary Fig. 1), suggesting that HAR genes play a specific role in differentiating higher-order cognitive networks from more primary/unimodal systems, with the highest differentiating expression of HAR genes in the DMN. Examining the other six functional networks, separately, did not show a significant enhancement of HAR gene expressions (Supplementary Table 4).

### HAR-BRAIN gene expression

With the set of HAR genes including genes involved in all sorts of functions across the entire human body (and thus not specific to ‘brain’), we continued by examining whether and if so how HAR genes that are related to brain processes may have played a specific role in the high evolutionary cortical expansion of cognitive functional networks in the human brain. We identified genes commonly expressed in brain areas using the GTEx database (https://www.gtexportal.org/), selecting 2,979 genes significantly more expressed in brain tissues compared to other available body sites (*q* < 0.05; FDR correction; we further refer to this set as BRAIN genes). As expected^23^, a large percentage of the total set of HAR genes were found to be highly expressed in brain tissues; 415 genes (24.3%) out of the full set of 1,711 HAR genes were observed to be significantly expressed in brain, a set from now on referred to as HAR-BRAIN genes (in contrast to HAR-nonBRAIN genes; Supplementary Table 3).

We then wanted to examine 1) whether the subset of HAR-BRAIN genes was more expressed particularly in regions of higher-order cognitive networks compared to the total set of HAR genes, and 2) to what extent HAR-BRAIN genes were more expressed in regions of higher-order cognitive networks, more than an average set of genes related to general brain processes (i.e., BRAIN genes). First, the cortical expression pattern of HAR-BRAIN genes significantly correlated with the pattern of human cortical expansion (*r*(53) = 0.486, *p* < 0.001; Fig. 3c). Furthermore, HAR-BRAIN genes showed significantly higher expression levels in regions of cognitive networks as compared to the SMN/VN (*t*(44) = 5.136, *p* < 0.001, FDR corrected; Fig. 3e), with the highest expression levels observed in cortical regions of the DMN (*t*(55) = 3.267, *p* = 0.002, FDR corrected, DMN versus the rest of the cortex; Fig. 3f). These effects were respectively ×3.2 and ×2.7 larger than the effect obtained by HAR-nonBRAIN genes (*t*(55) = 1.028, *p* = 0.309 and *t*(55) = 1.212, *p* = 0.232, separately). Notably, examining the other six functional networks, separately, did not show significant elevations of HAR-BRAIN gene expressions (Supplementary Table 5).

**Figure 3.**
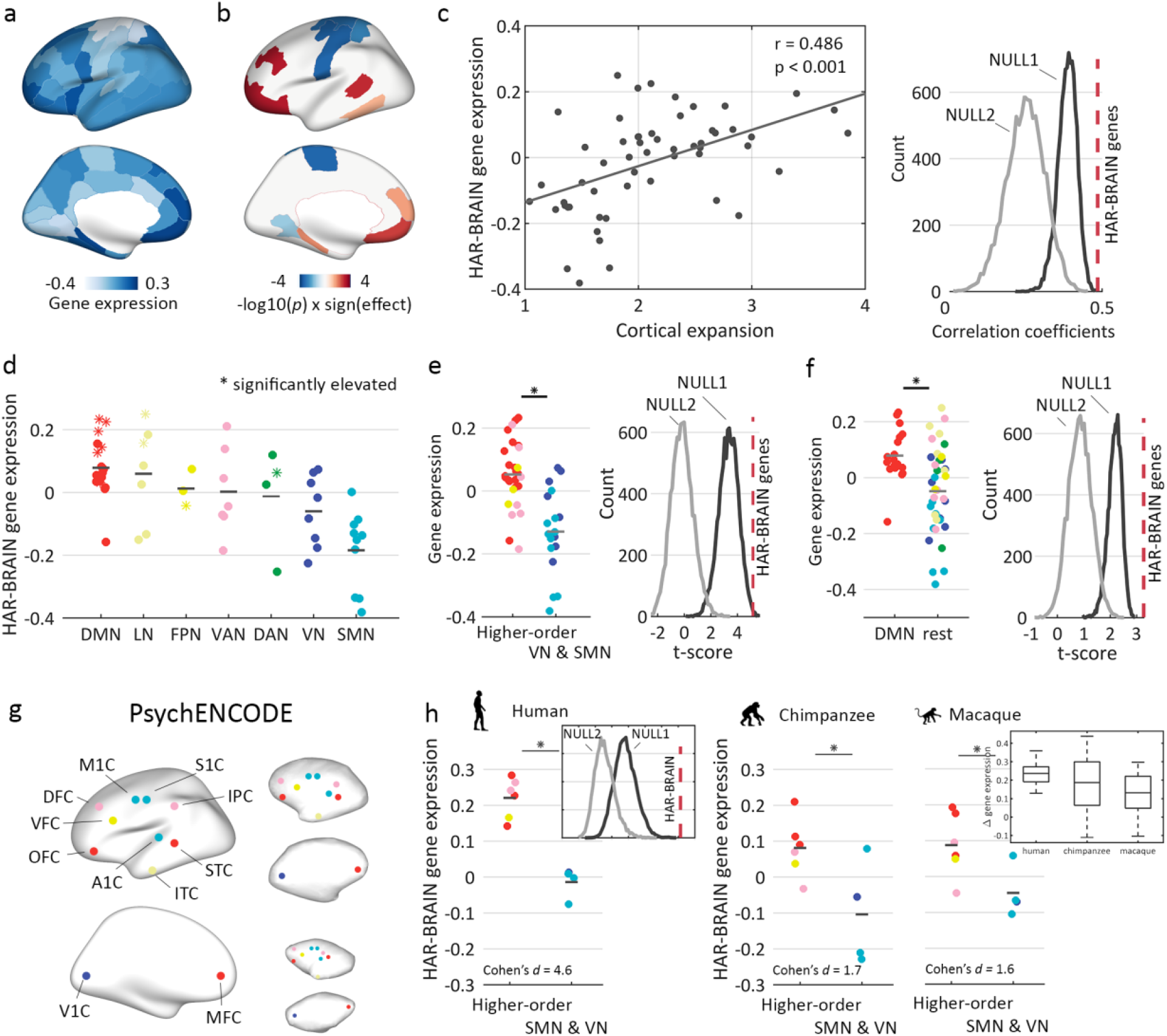
HAR-BRAIN gene expression. (a) Cortical gene expression of HAR-BRAIN genes. (b) Cortical map of the significance level of each cortical region obtained by permutation tests comparing expression of HAR-BRAIN genes to equally sized random gene sets taken from the total collection of BRAIN and ECE genes. (c) Association between gene expression profiles of HAR-BRAIN genes and cortical expansion between human and chimpanzee (left). The correlation coefficient is significantly higher than obtained by the null distribution generated by either BRAIN genes (NULL1) or ECE genes (NULL2; both *p* < 0.001, two-sided). (d) HAR-BRAIN gene expression within each of the seven functional networks ranked in a descending order according to their mean expression per network. DMN showed the highest HAR-BRAIN expression. ‘*’ indicates significant regions as in (b), with 6 out of the 10 significant regions (*p* = 0.008) present in the DMN. (e) HAR-BRAIN gene expressions in cognitive networks (DMN, FPN, and VAN) versus the SMN and VN (left), with permutation results demonstrated in the right panel (one-sided *p* = 0.003 and *p* < 0.001 for NULL1 and NULL2, separately). (f) HAR-BRAIN gene expressions in DMN versus the rest of the cortex (left), with permutation results demonstrated in the right panel (two-sided *p* < 0.001 for both NULL1 and NULL2). (g) PsychENCODE validation and comparative analysis. Matching brain areas as presented in the PsychENCODE dataset for the human (left), chimpanzee (upper right panel) and macaque (lower right panel). (h) Expression levels of HAR-BRAIN genes in regions of higher-order networks compared to areas of the SMN/VN in humans (*p* < 0.001), with weaker effects in chimpanzees and macaques. The largest differences in gene expressions are shown in humans, then chimpanzees, with the smallest in macaques (*p* = 0.002). ‘*’ indicates two-sided *p* < 0.05, FDR corrected. Central marks denote the mean gene expressions. Colors indicate the assignment of functional networks, as in Fig. 2b. M1C: primary motor cortex; S1C: primary sensory cortex; IPC: inferior parietal cortex; STC: superior temporal cortex; ITC: inferior temporal cortex; A1C: primary auditory cortex; OFC: orbital frontal cortex; VFC: ventral frontal cortex; DFC: dorsal frontal cortex; V1C; primary visual cortex; MFC: medial frontal cortex.

Second, we compared the level of expression difference of HAR-BRAIN genes between higher-order cognitive networks and SMN/VN using two types of null-distributions of expression differences generated by randomly selecting the same number of genes (i.e., 415) from the pool of 2,979 BRAIN genes (referred to as NULL1) and 8,686 ECE genes (referred to as NULL2). The elevated expression of HAR-BRAIN genes in regions of higher-order cognitive networks was significantly larger than both null distributions (*p* = 0.006 and *p* < 0.001 for NULL1 and NULL2, respectively; 10,000 permutations; Fig. 3e). The same result was also observed when examining DMN regions specifically (*p* < 0.001 for both NULL1 and NULL2; 10,000 permutations; Fig. 3f), suggesting a specific role of HAR-BRAIN genes in differentiating DMN regions from the rest of the brain. Permutation testing based on randomly shuffling cortical areas showed similar results (Supplementary Figure 5).

Further, we examined HAR-BRAIN expression for each of the cortical regions of the 7 functional networks separately; 10 cortical areas showed significantly high expression of HAR-BRAIN genes compared to a random selection of genes out of both BRAIN (NULL1) and ECE genes (NULL2) (FDR corrected, *q* < 0.05; Fig. 3b). Importantly, 7 out of these 10 regions described regions of the higher cognitive networks and 6 out of these 7 regions described DMN regions (which is beyond chance level of randomly selecting 10 regions out of 57, *p* = 0.034 and 0.008 for obtaining 7 higher-order network regions and 6 DMN regions, respectively; hypergeometric test). Together, these findings support the hypothesis of a specific role of HAR-BRAIN genes in the architecture of cognitive functional brain networks, above and beyond effects of general evolutionary conserved genes and general BRAIN genes.

### Chimpanzee-human comparative gene expression

Our analyses so far suggest that genes important for human brain evolution (i.e., HAR-BRAIN) are differentially more expressed in expanded regions of associative cognitive networks, in particular the DMN, compared with primary/unimodal areas. These findings do not yet provide direct information on whether HAR-BRAIN genes are *differently* expressed (or more specific, up-regulated) in the human brain compared to that of other primate species. To examine this, we used gene expression data from the PsychENCODE database (http://evolution.psychencode.org/)^24^, which describes gene expression of 11 comparable cortical regions in humans, chimpanzees, and macaques. Due to the lower spatial sampling of cortical regions (data of 3 DMN regions available), we limited our examination to a comparison between regions of cognitive networks and somatosensory/visual networks. First, we replicated the high expression of HAR-BRAIN genes in regions of higher-order cognitive networks compared to regions of the SMN/VN in humans (*t*(8) = 7.135, *p* < 0.001, FDR corrected; Fig. 3g), with this effect significantly exceeding both NULL1 and NULL2 (both *p* < 0.001, 10,000 permutations), confirming our AHBA-based findings of high HAR-BRAIN gene expression to go beyond expression levels of ECE and BRAIN genes in cognitive networks in the human brain. Second, the differentiating level of HAR-BRAIN gene expression between cognitive and primary/unimodal areas observed in humans (Cohen’s *d* = 4.605) was found to be ×2.7 larger than the effects found in chimpanzees (Cohen’s *d* = 1.695) and ×2.8 larger compared to macaques (Cohen’s *d* = 1.616). Chimpanzees and macaques showed only marginal higher HAR-BRAIN gene expression in regions of higher-order cognitive networks as compared to the SMN/VN (*t*(8) = 2.626, *p* = 0.030 for chimpanzees; *t*(8) = 2.504, *p* = 0.037 for chimpanzees). Statistical evaluation of this comparative effect across the three species showed a significant decreasing step-wise relationship of differences in HAR-BRAIN gene expression between regions of higher-order networks and the SMN/VN from humans (highest) to chimpanzees and macaques (lowest differentiating expression, Jonckheere-Terpstra test, *p* = 0.002). These findings expand the observations of significantly high HAR-BRAIN expression in cognitive network regions, now further showing that humans show up-regulated expression of HAR-BRAIN genes in brain areas involved in cognitive brain functions compared to chimpanzees.

### GWAS on DMN functional connectivity

We also wanted to examine the role of HAR and HAR-BRAIN genes in the inter-subject variation of default mode functional connectivity (DMN-FC) in today’s human population. We performed a GWAS (see methods) on 6,849 participants from the UK Biobank (February 2017 release)^25^ with DMN-FC as the phenotype of interest (see Methods). GWAS results for all single-nucleotide polymorphisms (SNPs) with minor allele frequency (MAF) > 0.0073 (i.e., minor allele count, MAC > 100^25^; to control unreliable results led by rare alleles) were annotated using the web-based platform FUMA (see Methods). We observed 24 independent (*r*^2^ < 0.1) genome-wide significant SNPs (*p* < 5 × 10^−8^; Fig. 4a) across 22 genomic loci. We functionally annotated 168 SNPs that were in high linkage disequilibrium (*r*^2^ ≥ 0.6) with one of these 24 independent SNPs and showed an association *p*-value of at least *p* < 1 × 10^−5^, and mapped these SNPs to genes using positional mapping, eQTL mapping, and chromatin interaction mapping (see Supplementary Methods)^26^. In total, this resulted in the observation of 195 genes to relate to DMN functional connectivity (Supplementary Table 6; 12 out of these 195 genes further annotated using brain-related eQTL and Hi-C mappings). Hypergeometric enrichment testing^26^ showed significant enrichment of the 195 genes in GWAS catalog reported gene-sets of brain-related disorders such as *bipolar disorder* (*p* = 7.35 × 10^−9^), *autism spectrum disorder, schizophrenia* (*p* = 1.93 × 10^−5^), *depressive symptoms* (*p* = 9.31 × 10^−5^), and *Alzheimer’s disease biomarkers* (*p* = 9.56 × 10^−4^). Moreover, these 195 genes showed enrichment for *intelligence* (*p* = 0.001) (see Fig. 4b and Supplementary Table 7 for a complete list). Furthermore, the set of 195 genes was significantly over-represented by HAR genes (*p* = 2.96 × 10^−4^; see a full list of 27 HAR genes in Supplementary Table 6; 6 HAR-BRAIN genes, *p* = 0.054), and these gene sets were found to significantly share similar pathways (permutation testing, *p* < 0.001, 10,000 permutations; Fig. 4c). These findings suggest that human DMN-FC relevant genes show overlap with gene-sets associated with several mental brain disorders and general human intelligence as well as show overlap with pathways related to evolutionary HAR genes.

**Figure 4.**
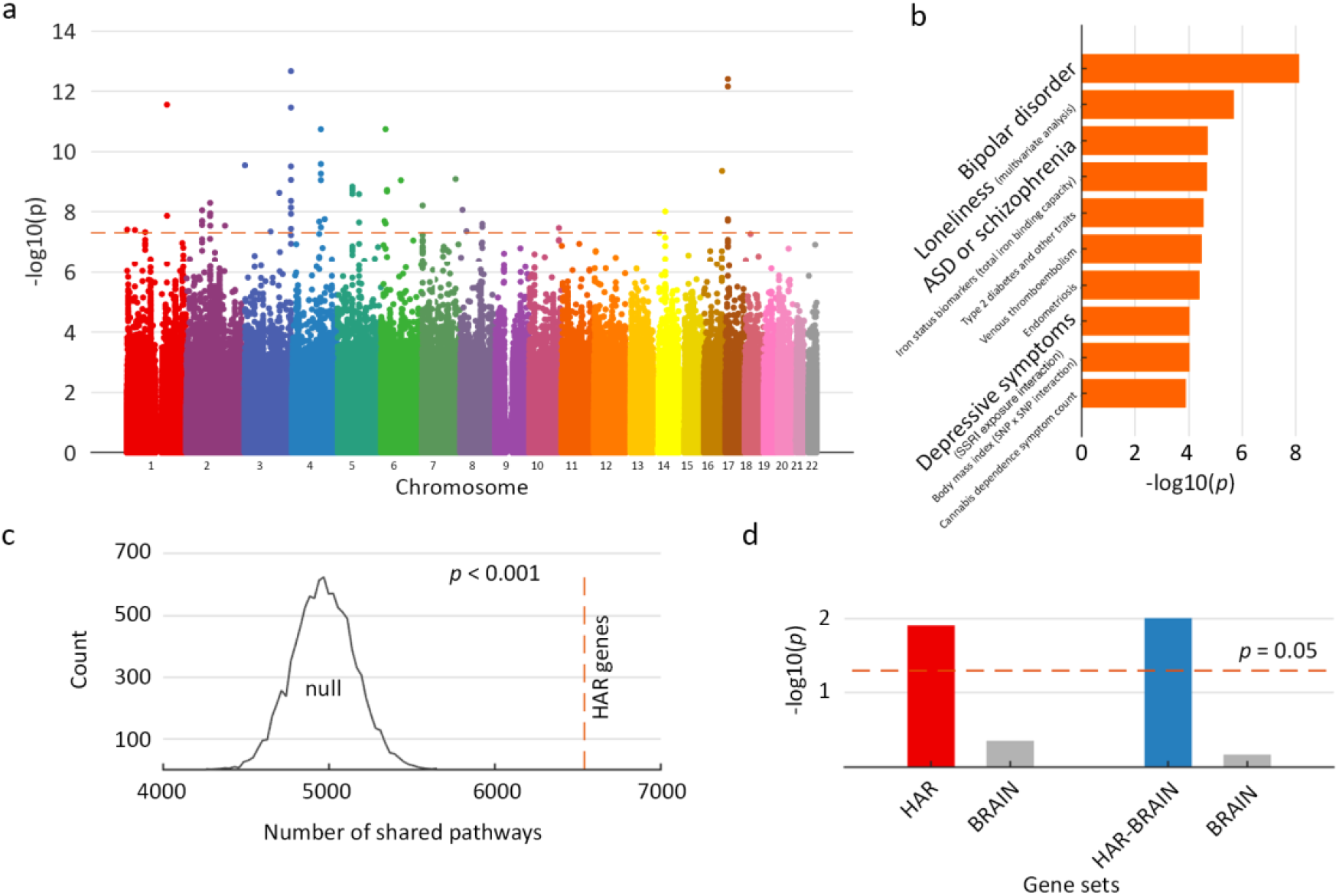
GWAS of DMN-FC. (a) Manhattan plot showing −log_10_-transformed two-tailed *p* value of each SNP from the GWAS analysis on the y axis and base-pair positions along the chromosomes on the x axis. Dotted red line indicates Bonferroni-corrected genome-wide significance (*p* < 5 × 10^−8^). (b) Enrichment of genes related to DMN-FC in GWAS catalog gene-sets (top 10 significant terms). (c) 2,288 pairs (within a total of 7,246 possible pathways) of DMN-FC risk genes and HAR genes were found to share pathways, significantly exceeding the null condition of DMN-FC risk genes and random genes (see Supplementary Methods for details). (d) MAGMA conditional gene-set analysis. The sets of HAR and HAR-BRAIN are associated with DMN functional connectivity conditional upon BRAIN genes. Dashed line indicates *p* = 0.05.

We further investigated the potential role of HAR genes in human DMN-FC by means of a MAGMA linear regression gene-set analysis^27^, which allows for a follow-up specific examination of the extent to which a certain set of genes (e.g. HAR/HAR-BRAIN genes) is associated with a phenotype (e.g., DMN-FC). First, gene-set analysis of the DMN-FC GWAS association results revealed the sets of HAR and HAR-BRAIN genes to be significantly associated with DMN-FC (standardized regression coefficient *β* = 0.016, *p* = 0.012 for HAR genes; *β* = 0.016, *p* = 0.010 for HAR-BRAIN genes; FDR corrected; Supplementary Table 8). An additional conditional gene-set analysis^27^, which included the set of BRAIN genes as a covariate in the linear regression model to examine the additive contribution of HAR genes above BRAIN genes, did show a significant effect of the association of HAR and HAR-BRAIN genes with DMN functional connectivity (*β* = 0.016, *p* = 0.012 and *β* = 0.017 *p* = 0.010, separately; Fig. 4d), which further supports the notion of HAR-BRAIN genes to be involved in inter-subject variation in DMN organization. SNP heritability enrichment analysis did unfortunately not result in further findings of significant enrichment of HAR in genetic variations in DMN-FC, which is likely related to HARs occupying relative very small regions in the whole genome (with a mean length of only 256 base pairs), reducing statistical power.

### Top strongest differentiating DMN genes

We continued by focusing on the evolutionary and biological properties of genes showing the highest levels of expression in DMN regions out of all 20,734 genes. For each of the 20,734 genes annotated in AHBA, we computed the level of over-expression in regions of the DMN by calculating the *t*-score for expression levels of the selected gene in regions of the DMN against the rest of the brain. The top 200 expressed genes (i.e. the set of genes showing the highest positive *t*-scores, referred to as DMN genes; all *p* < 0.004) were taken as the DMN’s most differentiating genes (Supplementary Table 9). Out of the top 200 DMN genes, we identified 37 to be HAR genes, particularly including 18 HAR-BRAIN genes, which greatly exceeded the chance level of randomly selecting 37/18 out of 20,734 (both *p* < 0.001, hypergeometric test, FDR corrected). As for an alternative selection of selecting the top 200, we also examined the top 53 genes based on *p* < 0.0014 (partial Bonferroni corrected) and top 469 genes with *p* < 0.01 (not corrected), which revealed comparable findings, see Supplementary Methods and Results.

To investigate whether the observed effect was restricted to the DMN or whether it represented a common brain effect, an additional permutation analysis was performed by shuffling region labels across the seven functional networks and re-computing the top genes for each of these random network assignments. We found the ratio of HAR genes in the set of top DMN genes to be significantly higher than the null condition (*p* < 0.001, 10,000 permutations), which confirmed a dominant role of HAR genes in DMN organization. To further examine the potential DMN specificity, we also selected the top 200 genes showing the largest differentiating gene expression in each of the other functional networks compared to the SMN/VN. In contrast to the DMN (revealing 18 overlapping genes with the set of HAR-BRAIN genes), the top 200 gene sets identified by the VAN, DAN, FPN, and LN, comprised respectively 12,8,7 and 7 HAR-BRAIN genes. However, only 2 out the top 200 DMN genes (CALB1, SEC24D) overlapped with the DMN-FC genes resulting from the GWAS results, which was not a significant effect (*p* = 0.223). Gene-set analysis on the set of DMN genes using hypergeometric testing as implemented in FUMA^26^ further showed significant over-representation of DMN genes in Gene Ontology (GO) terms related to cellular components of *dendrite* (*p* = 2.60 × 10^−5^), *somatodendritic compartment* (*p* = 4.49 × 10^−5^), *synapse* (*p* = 2.15 × 10^−4^), *perikaryon* (*p* = 7.45 × 10^−5^) and *neuron projection terminus* (*p* = 2.26 × 10^−4^), as well as molecular functions of *neuropeptide hormone activity* and *calcium activated potassium/cation channel activity* (*p* = 1.66 × 10^−5^ and 1.32 × 10^−4^; FDR corrected; Supplementary Table 10).

### HAR and DMN genes, cognitive abilities, social behavior, and psychiatric disorders

HAR/HAR-BRAIN and DMN genes were cross-referenced with the results of a recent large GWAS meta-analysis on intelligence performed on 269,867 individuals^28^. Competitive gene-set analysis^27^ revealed the sets of HAR/HAR-BRAIN genes to be significantly associated with individual variations in intelligence (*β* = 0.058, *p* = 5.12 × 10^−10^ for HAR genes; *β* = 0.076, *p* = 7.52 × 10^−16^ for HAR-BRAIN genes; Supplementary Table 8). Using conditional gene-set analysis^27^ with the general set of BRAIN genes taken as a covariate showed similar results (*β* = 0.056, *p* = 1.41 × 10^−9^ for HAR genes; *β* = 0.061, *p* = 3.17 × 10^−10^ for HAR-BRAIN genes). Intelligence was not found to be specifically associated with the set of DMN genes (*β* = 0.011, *p* = 0.078, n.s.), but a significant effect was observed for the subset of 37 intersected HAR-DMN genes (*β* = 0.022, *p* = 0.009).

We also linked HAR and DMN genes to genetic effects on human social behavior, which is thought to be more sophisticated in humans than that of other primate species^16^. GWAS summary statistics of the social behavior trait “Frequency of friend/family visits” based on a GWAS analysis on 383,941 individuals in the UK Biobank were obtained from the GWAS ATLAS web tool (http://atlas.ctglab.nl, ID 3216). HAR/HAR-BRAIN genes were found to be significantly associated with “Frequency of friend/family visits” (HAR genes: *β* = 0.037, *p* = 1.24 × 10^−7^; HAR-BRAIN genes: *β* = 0.039, *p* = 1.01 × 10^−7^; Supplementary Table 8), with effects unrelated to the set of BRAIN genes (conditional gene-set analysis, *β* = 0.037, *p* = 1.73 × 10^−7^ for HAR genes; *β* = 0.036, *p* = 2.63 × 10^−6^ for HAR-BRAIN genes). Furthermore, the DMN genes also showed a significant association with individual variation in this phenotype (*β* = 0.014, *p* = 0.023).

We further examined the potential association of HAR (and more specifically HAR-BRAIN) and DMN genes with schizophrenia, a disorder hypothesized to relate to human brain evolution^29, 30^. We used the SNP-based association statistics of a GWAS in 33,426 schizophrenia patients and 54,065 healthy controls^31^ provided by the Psychiatric Genomics Consortium (http://www.med.unc.edu/pgc/). Using gene-set analysis, we observed HAR/HAR-BRAIN genes to be significantly associated with genetic variants in schizophrenia (*β* = 0.056, *p* = 1.78 × 10^−11^ for HAR genes; *β* = 0.076, *p* = 1.82 × 10^−19^ for HAR-BRAIN genes; Supplementary Table 8). These results remained significant in the conditional gene-set analysis with BRAIN genes taken as a covariate (*β* = 0.018, *p* = 0.018 for HAR genes; *β* = 0.027, *p* = 0.001 for HAR-BRAIN genes). DMN genes were also found to be significantly enriched for genetic variants related to schizophrenia (*β* = 0.014, *p* = 0.028).

In addition to common variations indicated by GWAS, we alternatively examined the enrichment of HAR/HAR-BRAIN and DMN genes in rare variations of brain disorders using the NPdenovo database (http://www.wzgenomics.cn/NPdenovo/)^32^. Hypergeometric testing showed HAR and HAR-BRAIN genes to be significantly enriched in risk genes of ASD (*p* < 0.001 and *p* = 0.005, separately; FDR corrected) and schizophrenia (*p* < 0.001 and *p* = 0.008, separately; FDR corrected). DMN genes also showed significant enrichment in risk genes of ASD (*p* = 0.004, FDR corrected), but not schizophrenia (*p* = 0.264; see Supplementary Fig. 8).

Finally, we examined the potential association of evolutionary HAR/HAR-BRAIN and DMN genes with general brain changes related to psychiatric disorders. We used data from voxel-based morphometry (VBM) studies of cortical volume changes in five psychiatric brain disorders (schizophrenia, bipolar disorder, ASD, major depressive disorder [MDD] and obsessive-compulsive disorder [OCD]; see Methods) and created a cortical map describing the distribution of cortical changes to brain regions across these five psychiatric disorders (including in total meta-data of 260 VBM studies, see Supplementary Methods). The spatial pattern of disorder involvement across the cortex was significantly associated with the gene expression pattern of HAR-BRAIN genes (*r*(55) = 0.437, *p* < 0.001, with the cortical volume controlled; Fig. 5b; *r*(55) = 0.221, *p* = 0.098 for HAR genes), an effect significantly exceeding the effect obtained by general BRAIN genes (i.e., NULL1; *p* = 0.022, 10,000 permutations) and ECE genes (i.e., NULL2; *p* < 0.001, 10,000 permutations). For an out-group analysis, the cortical pattern of HAR-BRAIN gene expression did not correlate to the disease map of five alternative, non-psychiatric disorders (amyotrophic lateral sclerosis, stroke, alcoholism, insomnia, fibromyalgia; *r* = 0.141, *p* = 0.302).

**Figure 5.**
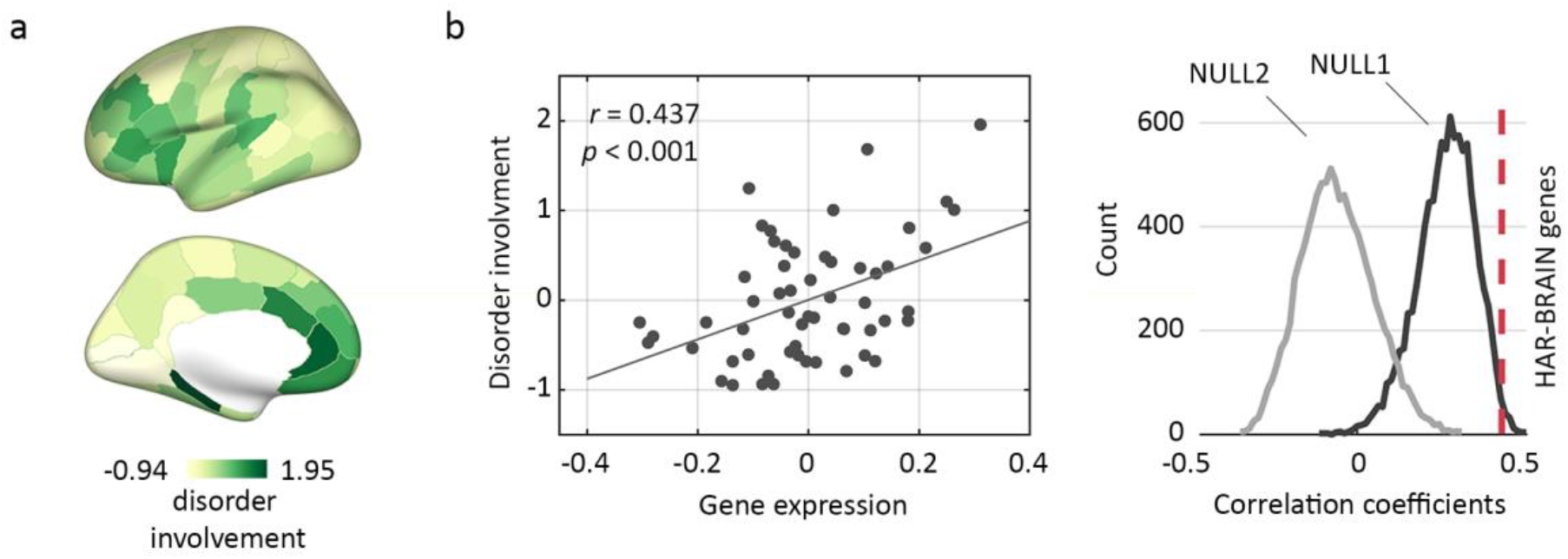
Cortical disorder involvement and HAR-BRAIN gene expression. (a) Brain maps of cortical involvement across five major psychiatric disorders (e.g., schizophrenia, bipolar disorder, autism spectrum disorder, major depression, and obsessive-compulsive disorder). (b) Correlation between HAR-BRAIN gene expression and disorder involvement (*r* = 0.437, *p* < 0.001; corrected for cortical volume), and comparisons of the effect to null distributions generated by random BRAIN (NULL1; *p* = 0.022) and ECE genes (NULL2; *p* < 0.001).

## Discussion

Our comparative neuroimaging and genetic findings together suggest that recent evolutionary changes in the human genome have played a central function in the expansion and cortical organization of cognitive functional networks in the human brain, potentially in service of evolutionary specialization of higher-order cognitive and social brain function.

Our results suggest high levels of cortical expansion in regions of both the FPN and DMN in humans. This is compatible with prior observations of cortical variation between macaque and human^33^, showing large expansion of associative prefrontal, temporal, and parietal areas in humans^33, 34^. Our chimpanzee-human comparison now further demonstrates a relatively large expansion of the lingual gyrus in humans, which might attribute to humans’ largely evolved functions of word processing and language compared to chimpanzees^35^. The evolutionary expansion pattern has been suggested to be comparable to the pattern of cortical variation in today’s human population^36^, showing larger brains to potentially display relative larger multi-modal associative areas, effects linked to individual variation in cognitive abilities^36^. The current observation of high expression of HAR genes in brain areas of cognitive networks may suggest that genes linked to hominization had a central role in the process of cortical development of these multimodal association areas.

Our findings indicate that the expression of genes related to human evolution (HAR genes) and in particular genes related to human brain evolution (HAR-BRAIN genes) are not evenly expressed across cortical areas but instead, are more expressed in areas related to higher-order cognitive processing. HAR genes, representing conserved loci with elevated divergence in humans^17, 19^, have been argued to function as important neuronal enhancers^37^ and to be key players in biological processes of nervous system development and neurogenesis, amongst other (Supplementary Table 11). The high expression of HAR genes in regions of central cognitive networks, and most pronounced in areas of the DMN, may thus reflect an enhanced neuronal complexity in these areas of the human brain^38, 39, 40^. Such regional variation in gene expression is potentially associated with distinct cytoarchitectural structural types in the cortex, where regions of higher-order cognitive networks display a different cytoarchitectonic organization compared to unimodal brain areas^41^. This is also consistent with findings of association areas in the human brain^42^ to show transcriptional profiles enriched for genes specific to the organization of supragranular layers. The spatial layout of functional networks in humans has been shown to be nicely captured by coupled transcription profiles of genes enriched in supragranular layers^43^ and genes related to ion channel activity and synaptic function^44^. These observations are further in line with the notion of genes related to the resting-state brain activity of the DMN to have greater expression in neurons^9^ and corroborate on the current results of genes with high transcriptional differentiation in the DMN to display broad involvement in the formation of synapses and dendrites. Moreover, our observation of up-regulated HAR-BRAIN gene expression in cognitive networks in humans compared to chimpanzees and macaques implicates an evolutionary enhanced complexity of neuronal connectivity in cognitive networks. This might be further related to that humans have potentially a longer period of neuronal progenitor expansion compared with chimpanzees and macaques contributing to a differentiated neuronal number and cortical size^45^.

Our findings show that genes highly expressed in the DMN contain genetic variants related to human intellectual ability and social cognition and behavior, further implicating a shared genetic origin between, and a potential role of, DMN organization and intellectual abilities and social cognition. This is compatible with neuroimaging twin studies showing a genetic correlation between IQ and gray matter morphology of DMN regions like the medial frontal cortex and parahippocampal gyrus^46^. The DMN has indeed been argued to be central to human self-projection abilities that include planning the future^47^, remembering the past, theory of mind, and navigation^13^, of which humans show a higher complexity as compared to chimpanzees^15, 48^. This central cognitive system has also been noted to comprise highly connected network hubs like the precuneus and inferior parietal lobule^49^, regions thought to be involved in multimodal information integration^50^, a key aspect of high order cognitive and social brain function. The combined findings of enrichment of HAR genes in DMN regions and their role in functional connectivity and variation in intelligence in the human population suggest a central role for HAR genes in the evolution of the human brain cognitive networks in the period following the divergence of the human and chimpanzee lineage.

We suggest that some of the genes found at the intersection of HAR, BRAIN and DMN genes (Fig. 1) directly relate to the development of the human central nervous system. For example, CDH8, CDH9, and CDH10, encoding a type II classical cadherin, are involved in synaptic adhesion, axon outgrowth and guidance^51^, and are known to be involved in several psychiatric disorders like autism^51^. CBLN1 encodes a cerebellum-specific precursor protein, precerebellin, and is important for synapse integrity and synaptic plasticity together with NRXN1 and GRID2^52^. CA10, which belongs to the carbonic anhydrase family of zinc metalloenzymes, is believed to be central in the development of the central neural system by coordinating neurexins, which are presynaptic cell-adhesion molecules that bind to diverse postsynaptic ligands, and are linked to neuropsychiatric disorders^53^. KCNB2 regulates potassium voltage-gated channel and is known to be essential in regulating neurotransmitter release and neuronal excitability^54^.

Our findings also point into the direction of genes enriched in cognitive functional networks in humans to be involved in the development of psychiatric disorders. The cortical expression patterns of HAR-BRAIN and DMN genes show significant overlap with the pattern of cortical involvement across five mental brain disorders, with large involvement observed in lateral and medial prefrontal cortices, key ‘brain hubs’ and components of high-order networks and identified to be generally implicated in the anatomy of a wide range of brain disorders^55, 56^. These brain regions also show a hyper-expansion of surface area in the developing brain of autistic children^57^, which is in line with our findings of associated HAR gene expression and evolutionary cortical expansion. Furthermore, HAR and DMN genes are observed to be significantly associated with the genetic architecture for schizophrenia and ASD, disorders that are often reported to involve disturbed DMN functional connectivity^58^. These new findings are consistent with reported genetic associations between the DMN and psychiatric disorders^59^ and support the notion of genes related to evolutionary adaptations and brain development to potentially contribute to default mode alterations in brain disorders^59^.

Several methodological points have to be discussed when interpreting the findings of our study. We used predefined population-based resting-state functional networks to categorize cortical regions into seven functional networks to link data from distinct modalities. Several other spatial variations of these networks have been suggested^60^ (see Supplementary Results for validation of our findings with alternative functional network definitions, including variations of the DMN) and these network divisions may overlook functional heterogeneity across cortical regions and participation of brain regions in multiple networks. For example, the orbital frontal cortex, in the used network parcellation included in the LN, showed highest HAR-BRAIN expression in this network, and has been thought to be important in multiple cognitive functions such as learning, prediction, decision making for emotional and reward-related behaviors, and subjective hedonic processing^61^ and is often also included in the DMN^6^. Second, the set of HAR genes as used in this study was selected ‘as-is’ from the study of Doan et al.^19^, which defined HAR segments of the human genome from recent studies comparing humans and other species. HARs-associated genes were labeled as those where HARs are within the introns, within or near (less than 1kb) 5’ and 3’ UTRs, or are the closest flanking gene that was less than 2.1mb away, (with 70% being less than 500kb away)^19^. Other mapping approaches can be used to identify and further specify HAR gene sets linked to specific functions, for example by using Hi-C and eQTL. We examined alternative sets of 196 genes mapped from HAR regions using brain-related Hi-C and eQTL datasets from the PsychENCODE Consortium^62^, and a set of 396 genes related to ASD-linked HAR mutations identified using massively parallel reporter assays (MPRA)^19^. These alternative selections and allocations of HAR genes revealed highly consistent findings (data presented in detail in Supplementary Results).

Our findings implicate that evolutionarily accelerated regions in the human genome have played a central role in the expansion and organization of cognitive functional networks in the human brain. Examining the role of genes important for human brain evolution and their role in the development of functional brain architecture provide a new window for examining the biological mechanisms of human brain disorders.

## Methods

### Cortical expansion

In vivo MRI data from 29 adult chimpanzees and 50 adult human subjects were analyzed (see Supplementary Methods for details). T1-weighted scans of both species were processed using FreeSurfer for tissue classification, cortical ribbon reconstruction, and brain parcellation^63^. Pial surface reconstructions were used for vertex-to-vertex mapping across chimpanzee and humans and subsequent computation of vertex- and region-wise expansion. Validation analysis was performed using the chimpanzee-human BB38 atlas that describes homologous cortical regions between two species^29, 64^ (Supplementary Methods).

### Gene expression data

#### AHBA

Cortical gene expression patterns were taken from the transcriptomic data of the Allen Human Brain Atlas (AHBA, http://human.brain-map.org/static/download), including a highly detailed dataset of microarray gene expression data from brain samples of six human donors (all without a history of neuropsychiatric or neuropathological disorders, demographics tabulated in Supplementary Table 12). Data included expression levels of 20,734 genes represented by 58,692 probes for each cortical region of the left hemisphere^65^. Tissue samples were spatially mapped to each of the cortical regions of the FreeSurfer Desikan-Killiany cortical atlas (114 cortical regions, 57 per hemisphere, DK-114)^66, 67^, based on their distance to the nearest voxel within the cortical ribbon of MNI 152 template (and BB38 atlas for validation, see Supplementary Results). Samples were normalized to Z scores and averaged across regions (see Supplementary Methods), resulting in a subject × region × gene (6 × 57 × 20,734) data matrix.

#### PsychENCODE

Cortical transcription data in adult humans, chimpanzees, and macaques were obtained from the PsychENCODE database (http://evolution.psychencode.org/)^24^. The PsychENCODE database provides gene expressions of 16,463 genes for 11 cortical locations in humans (6 subjects), chimpanzees (5 subjects) and macaques (5 subjects). As for the AHBA data, gene expression data were normalized to *Z* scores across cortical regions within each brain, resulting in gene expression matrices of the size of 6 × 11 × 16,463, 5 × 11 × 16,463, and 5 × 11 × 16,463 for human, chimpanzee, and macaque, respectively. Normalized gene expression data was averaged across individual brains to obtain a group level gene expression matrix of the size of 11 × 16,463 for each species.

### HAR genes

Genes located in human accelerated regions (HARs) of the human genome were taken as the list of genes presented by a comparative genome analysis representing genomic loci with accelerated divergence in humans^19^. A total number of 2,737 human accelerated regions were identified, representing a list of 2,143 HAR-associated genes^19^. 1,711 out of the 2,143 HAR genes were described in the AHBA dataset (referred to as HAR genes in the current study) with their gene expression profiles averaged across datasets.

### BRAIN genes

BRAIN genes were selected as the set of genes commonly expressed in human brain tissue using the Genotype-Tissue Expression (GTEx) database (data source: GTEx Analysis Release V6p; https://www.gtexportal.org/). The GTEx portal contains 56,238 gene expression profiles in 53 body sites collected from 7,333 postmortem samples in 449 individuals. From these 56,238 genes, a total number of 2,823 genes were identified as BRAIN genes showing significantly higher expressions in brain sites than non-brain sites (one-sided *t*-test and an FDR corrected *q* < 0.05 were used). 405 of these 2,823 genes overlapped with the set of HAR genes, referred to as HAR-BRAIN genes.

### GWAS on DMN-FC

GWAS analysis was performed based on combined genetic and MRI data of 6,849 participants from UK Biobank (http://www.ukbiobank.ac.uk). Individuals of European ancestry were included, and available anatomical T1 and resting-state fMRI (rs-fMRI) data were preprocessed (see Supplementary Methods). Functional connectivity (FC) was computed as the Pearson correlation coefficient between time-series of every pair of cortical regions in the DK-114 atlas^66, 67^. FC strength of the DMN (DMN-FC) was taken as the sum of FC between regions of the DMN divided by the sum of FC between non-DMN regions and used as our phenotype of interest. GWAS on DMN-FC was conducted in PLINK v1.90^68^, using an additive linear regression model, controlling for covariates of age, sex, twenty European-based ancestry principal components, genotyping array, and total brain volume (derived from the T1 image). Stringent quality control measures were applied to the summary statistics of the GWAS (see Supplementary Methods and ^28^ for a detailed description of the used procedures). SNP-based results were mapped and annotated using FUMA (Functional Mapping and Annotation of Genome-Wide Association Studies)^26^. MAGMA (Multimarker Analysis of GenoMic Annotation) gene-set analysis was used to examine the association of HAR genes with genetic variants of DMN-FC^27^.

### Top DMN genes

For each of the 20,734 AHBA genes, a two-sided two-sample *t*-test was performed between expression levels of regions of the DMN and regions of the other resting-state networks. Genes showing the top 200 largest *t*-scores (showing *p* < 0.004, uncorrected) were selected and referred to as DMN genes (consistent results were obtained for the set of genes reaching *p* < 0.05, corrected for multiple comparisons and alternatively the set of genes reaching *p* < 0.01 without correction; see Supplementary Results). The enrichment of HAR genes in top DMN genes was statistically evaluated using a hypergeometric test. Gene-set analysis was performed for the set of top DMN genes by means of the hypergeometric test implemented in GENE2FUNC in FUMA (http://fuma.ctglab.nl)^26^ (see Supplementary Methods for further details).

### Cortical involvement in psychiatric disorders

The BrainMap database was used to assess cortical involvement across five major psychiatric disorders (schizophrenia, bipolar disorder, ASD, MDD, and OCD, including in total 260 studies) (http://www.brainmap.org). Disease voxel-based morphometry (VBM) data of 260 case-control studies present in BrainMap were extracted using the Sleuth toolbox^69^ and meta-analyses were conducted for each disorder using the GingerALE toolbox^70^. Resulting brain maps of activation likelihood estimation (ALE) were registered to the MNI 152 template in the Freesurfer space and regional ALE was computed by averaging ALEs of all voxels within each cortical region of the DK-114 atlas. Regional averaged ALE was transformed to Z-score and then averaged into a cross-disorder cortical involvement map describing per region the level of involvement across the five major psychiatric disorders (see Supplementary Methods for details).

### Statistical analysis

Pearson’s correlation was used to examine the association of the profile of cortical gene expression with the pattern of chimpanzee-to-human cortical expansion. Two-tailed two-sample *t*-test was used to statistically test the difference in evolutionary cortical expansion and mean gene expression of HAR and HAR-BRAIN genes between regions of higher-order cognitive networks (e.g., the DMN, FPN, and VAN) and regions of the SMN and VN. Similar analysis was conducted between each of the functional network and the rest of the brain. Results reaching an FDR corrected *q* < 0.05 were taken as statistically significant. Permutation testing (10,000 permutations) was used to differentiate effects of HAR-BRAIN genes from effects of general BRAIN genes (referred to as NULL1) and genes associated with evolutionarily conserved elements of the human genome (ECE genes, referred to as NULL2). ECE genes were obtained from evolutionarily conserved elements in the human genome with length larger than 200 base pairs as described in ^22^ and mapped to genes when they fall inside the genomic location provided by the gene^27^, resulting in a set of 9,125 genes. For each permutation (for NULL1 and NULL2), 415 genes (the same size as the number of HAR-BRAIN genes) were randomly selected from the pool of 2,927 BRAIN genes or 9,125 ECE genes, separately, and the same effects (e.g., correlation or *t*-test) were computed for this random set to generate an empirical null-distribution (i.e., noted as the NULL1 distribution for BRAIN genes and NULL2 distribution for ECE genes). The original effects were assigned a two-sided p-value by comparison to the effects in the null-distributions, as the proportion of random permutations that exceeded the original score (of HAR-BRAIN genes).

## Acknowledgments

The work of M.P.v.d.H. was supported by an ALW open (ALWOP.179) and VIDI (452-16-015) grant from the Netherlands Organization for Scientific Research (NWO) and a Fellowship of MQ. Y.W. was supported by China Scholarship Council (201506040039). P.R.J. was supported by the Sophia Foundation for Scientific Research (SSWO, grant s14-27). L.L. was supported by National Institute of Mental Health (MH100029). D.P. was supported by The Netherlands Organization for Scientific Research (NWO VICI 453-14-005). Primate work was supported by National Institutes of Health Grants P01AG026423 and National Center for Research Resources P51RR165 (superseded by the Office of Research Infrastructure Programs/OD P51OD11132) to the Yerkes National Primate Research Center, and by the National Chimpanzee Brain Resource, R24NS092988. Human neuroimaging data was kindly provided by the Human Connectome Project, WU-Minn Consortium (Principal Investigators: David Van Essen and Kamil Ugurbil; 1U54MH091657) funded by the 16 NIH Institutes and Centers that support the NIH Blueprint for Neuroscience Research; and by the McDonnell Center for Systems Neuroscience at Washington University. The Genotype-Tissue Expression (GTEx) Project was supported by the Common Fund of the Office of the Director of the National Institutes of Health, and by NCI, NHGRI, NHLBI, NIDA, NIMH, and NINDS. The GTEx data used for the analyses described in this manuscript were obtained from: the GTEx Portal V6p release on 11/08/2017. The genetic analyses were carried out on the Genetic Cluster Computer, which is financed by the Netherlands Scientific Organization (NWO: 480-05-003), Vrije Universiteit, Amsterdam, The Netherlands, and the Dutch Brain Foundation, and is hosted by the Dutch National Computing and Networking Services SurfSARA. This research has been conducted using the UK Biobank resource under application number 16406. We would like to thank Mats Nagel (VU Amsterdam) and Ting Qi (Max Planck Institute for Human Cognitive and Brain Sciences) for helpful discussions and suggestions.

## Author Contributions

Y.W. and S.C.d.L. performed the analyses. M.P.v.d.H. conceived the idea of this study and supervised analyses. L.L., T.M.P., and J.K.R. collected chimpanzee MRI data. L.H.S. and D.J.A. contributed to chimpanzee data processing. P.R.J., J.E.S., and K.W. prepared genetic data. D.P. supervised the genetic analysis pipeline. Y.W. and M.P.v.d.H. wrote the manuscript with contributions from all coauthors.

## Data availability statement

The gene expression data that support the findings of this study are available in the Allen Brain Atlas (http://human.brain-map.org). The human MRI data (in the part of the cortical expansion) that support the findings of this study are available from the Human Connectome Project (https://www.humanconnectome.org). The chimpanzee MRI data that support the findings of this study are available from the corresponding author upon reasonable request. The T1-weighted MRI, resting-state fMRI, and genotype data (in the GWAS part) that support the findings of this study are available in the UK Biobank (https://www.ukbiobank.ac.uk). The GWAS summary statistics for “schizophrenia” that support the findings of this study are available from the Psychiatric Genomics Consortium (http://www.med.unc.edu/pgc). The GWAS summary statistics for “Frequency of friend/family visits’ that support the findings of this study are available from the GWAS ATLAS webtool (http://atlas.ctg.nl/traitDB/3216). The cross-disorder VBM data that support the findings of this study are available in the BrainMap (http://www.brainmap.org).

## Code availability

All code is available from the corresponding author upon reasonable request.

## Ethics statement

Data of chimpanzees were acquired under protocols approved by the YNPRC and the Emory University Institutional Animal Care and Use Committee (IACUC, approval #: YER2001206). Data of humans were acquired previously by external sources. All individuals included in the study provided informed consent. All studies were approved by local ethics committee for research in humans as described in original publications.

