## Supplementary Information for "Genetic correlates of evolutionary adaptations in cognitive functional brain networks and their relationship to human cognitive functioning and disease"

### Table of Contents

SUPPLEMENTARY METHODS 3

*MRI data* 3

*MRI processing* 4

*Cortical expansion* 5

*BB-38 chimpanzee-human atlas* 6

*Human transcription data from AHBA* 7

*HAR identification using brain-related Hi-C and eQTL* 9

*Functional networks* 9

*UK Biobank MRI and GWAS on DMN functional connectivity* 10

*Pathway-sharing analysis* 17

*Top genes differentiating genes of the default-mode network* 17

*Gene-set analysis based on GWAS of cognitive ability, social behavior, and schizophrenia* 18

*Cortical vulnerability in psychiatric disorders* 19

SUPPLEMENTARY RESULTS 21

*HAR genes identified by brain-related Hi-C and eQTL* 21

*HAR genes identified by MPRA* 22

*Top genes differentiating the default-mode network* 23

*Examination of the potential effects of tissue sample number* 24

*Alternative cortical parcellation atlases* 25

*Alternative functional network divisions* 26

*Directly mapping AHBA tissue samples to functional networks* 27

*Alternative parameters in processing* 29

*HAR-BRAIN gene expression in subcortical regions* 31

*The potential age and sex effect on expansion* 31

*Permutation testing of cortical expansion* 33

REFERENCES 33

SUPPLEMENTARY TABLES 38

Supplementary Table 2. Cortical expansion in each functional network 38

Supplementary Table 4. HAR gene expression in each functional network 38

Supplementary Table 5. HAR-BRAIN gene expression in each functional network 38
Supplementary Table 7. Enrichment of DMN-FC associated genes in GWAS catalog reported gene sets 38
Supplementary Table 8. Enrichment of HAR, HAR-BRAIN, BRAIN, and DMN genes in GWAS results 39
Supplementary Table 10. Over-representation of the top 200 DMN genes 40
Supplementary Table 11. Over-representation of HAR genes 41
Supplementary Table 12. Demographics of human donors included in the AHBA 42
Supplementary Table 13. Over-representation of the top 53 DMN genes 42
Supplementary Table 14. Over-representation of the top 469 DMN genes 44
SUPPLEMENTARY FIGURES 46

NOTE: Supplementary Tables S1, S3, S6, and S9 are in formats not supported by bioRxiv and available by request from the authors.

### Supplementary Methods

#### *MRI data*

We calculated cortical expansion from in vivo MRI data from 29 chimpanzees (*Pan troglodytes*; age: 30.2 ± 12.6, all female) and 30 humans (*Homo sapiens*; age: 29.8 ± 3.2 years; all female). Chimpanzees were accommodated at the Yerkes National Primate Research Center (YNPRC) in Atlanta, Georgia. All procedures were implemented under protocols approved by the YNPRC and the Emory University Institutional Animal Care and Use Committee (IACUC, approval #: YER-2001206). No new MRI data was acquired for this study. All chimpanzee MRIs were obtained from a data archive of scans obtained prior to the 2015 implementation of U.S. Fish and Wildlife Service and National Institutes of Health regulations governing research with chimpanzees. All the scans reported in this publication were completed by the end of 2012. All MRI scans are part of the National Chimpanzee Brain Resource (http://www.chimpanzeebrain.org).

Chimpanzee MRI scans were collected from chimpanzees under anesthesia with isoflurane (1%) (protocols described in detail in [1]). Chimpanzees were immobilized with ketamine (2-6 mg/kg). Constant observation was given by veterinary staff for chimpanzees before, during, and after scanning. Head motion was minimized using foam cushion and elastic straps. A standard circularly polarized (CP) birdcage coil was used because they did not fit the standard phase-array coil designed for humans. T1-weighted MPRAGE imaging of all chimpanzees was obtained on two Siemens 3T Trio Tim Scanners (Siemens Medical System, Malvern, PA) with the following parameters: slice thickness = 0.8 mm, voxel size = 0.8 × 0.8 × 0.8 $mm^3$, TR = 2,600 ms, TE = 3.06 ms, matrix size = 256 × 256 × 192, FOV = 224 × 224, flip angle = 8 degree, scanning time = 16 min.

Data of human subjects were randomly selected from the Q3 data release of the Human Connectome Project (HCP, http://www.humanconnectomeproject.org/)[2]. Human MRI scans were acquired on a Siemens Skyra 3T scanner (Siemens Medical System, Malvern, PA) with a customized SC72 gradient insert[2]. T1-weighted MPRAGE images were collected with following scanning parameters: voxel size = 0.7 × 0.7 × 0.7 $mm^3$, TR = 2400 ms, TE = 2.14 ms, matrix = 320, 256 sagittal slices, slice thickness = 0.7 mm, FOV = 224 × 224, flip angle = 8 degree, scanning time = 7:40 min.

***MRI processing***

Chimpanzee and human T1-weighted MRI data were processed in the FreeSurfer software[3, 4] for brain tissue segmentation and cortical mantle reconstruction of each subject. Chimpanzee-to-human cortical expansion was computed as follows, inspired by a recent human-chimpanzee morphological comparison based on FreeSurfer[5]. Individual reconstructed pial surfaces of both chimpanzees and humans were re-meshed to an identical number of vertices. FreeSurfer inflated the white matter – gray matter surface of each subject to a sphere, and registered the sphere to the standard reference ($FS_HOME/average/?h.average.curvature.filled.buckner40.tif) by aligning the curvature data, resulting in a surface file of the registered sphere. Next, for each vertex *i* (namely, a target vertex) on the registered sphere of the fsaverage, we extracted the face (i.e., a triangle formed by three vertices) comprising the location of vertex *i* on the registered sphere of each subject. To do this, we used Barycentric coordinates:

$$\boldsymbol{P}_i = [x_i, y_i, z_i] \qquad [1]$$

$$\begin{bmatrix} u \\ v \\ w \end{bmatrix} \times \begin{bmatrix} \boldsymbol{P}_1, 1 \\ \boldsymbol{P}_2, 1 \\ \boldsymbol{P}_3, 1 \end{bmatrix} = [\boldsymbol{P}_t, 1] \qquad [2]$$

where $\boldsymbol{P}_i$ is a vector of coordinates of a vertex *i*, $\boldsymbol{P}_1$, $\boldsymbol{P}_2$, $\boldsymbol{P}_3$ are coordinates of three vertices forming a face, $\boldsymbol{P}_t$ contains coordinates of the target vertex, and *u*, *v*, and *w* are numbers following $u + v + w = 1$. Given a target vertex on the registered sphere of the fsaverage template, we calculated *u*, *v*, *w* for each face on subjects' registered sphere and selected the face when all $u, v, w > 0$, which indicates that the location of the target vertex is comprised in the selected face. Using the calculated *u*, *v*, *w*, we generated coordinates of a new vertex within the selected face on subject's pial surface according to equation [2]. This way, pial surfaces of all chimpanzee and human subjects were re-meshed, resulting in spatially matched vertices across all subjects.

The re-meshed pial surfaces of all subjects were examined manually for both chimpanzee and human parcellations (i.e., 114- or 219-region subdivision of the Desikan Killiany atlas[6, 7]). An inconsistency of parcellation of the cuneus in the chimpanzees was noticed due to an inaccuracy in the registration process. Therefore, the corresponding annotation files were manually corrected for the parcellation of the cuneus lobe for all chimpanzees (according to the criteria that cuneus is bounded anteriorly by the parieto-occipital sulcus and inferiorly by the calcarine sulcus[8]). As FreeSurfer failed to properly parcellate the parahippocampal gyrus and entorhinal cortex in five of the 29 chimpanzees, we excluded these two regions in all cortical expansion-relevant analyses. Note that we also further validated the expansion and gene expression results by using the BB-38 homologous chimpanzee-human cortical atlas[9] (see below).

#### *Cortical expansion*

Cortical expansion was computed based on the reconstructed pial surfaces of both chimpanzees and humans. First, the area was calculated for each face within the re-meshed pial surface of each subject. A regional-level cortical area ($S_i$) was computed by summing up face areas within each cortical region, for all regions of the atlas (DK-114[6, 7]; also 219-region subdivision [DK-219] and the chimpanzee-human cytoarchitectonic cortical atlas [BB-38] for validation purposes, see below). Normalized cortical area was then obtained by dividing the regional area by the area of the whole cortex. Regional-level area was used to calculate cortical expansion between every pair of chimpanzee and human subjects by

$$E_{i,j} = \frac{S_{human,i} - S_{chimp,j}}{S_{chimp,j}} \qquad [3]$$

with $E_{i,j}$ denoting the expansion from chimpanzee $j$ to human $i$. They yielded a total of 870 (i.e., 29 × 30) chimpanzee-to-human expansion maps. For each expansion map, expansion levels were transformed to $Z$ scores across cortical regions. Then a one-sample $t$ test was performed for each region across 870 expansion maps to identify cortical regions with significant above- or below-average expansion levels. Finally, a group-wise region-wise cortical expansion map was made by averaging among the 870 chimpanzee-to-human comparisons.

***BB-38 chimpanzee-human atlas***

Results regarding the chimpanzee-human cortical expansion were validated using the BB-38 atlas that describes homologous cortical areas across the two species. In 1950, von Bonin and Bailey reported this cortical atlas[10], assessing cytoarchitectural homologies of the chimpanzee cortex based on the human cortical atlas of von Economo and Koskinas[11]. They provided detailed descriptions of 44 cytoarchitecturally distinct cortical regions of the chimpanzee brain, and how these regions relate to the human brain in terms of cytoarchitectural properties such as cortical layer thickness, neuronal cell type, size, and density[10]. Using a FreeSurfer version of the Von Economo - Koskinas atlas[8], we created the BB-38 chimpanzee-specific and homologous

BB-38 human-specific cortical atlas, with 6 of the original 44 regions (PCop, FCop, FDgamma, PA, PD, and TC) excluded because they could not be properly segmented in FreeSurfer (as they were too narrow, located within a sulcus, or embedded within another cortical region). Segmentation and atlas building followed the same procedures as described in prior literatures[8, 9].

***Human transcription data from AHBA***

Microarray gene expression dataset was collected from postmortem brains of six human donors and downloaded from the Allen Human Brain Atlas (AHBA) (http://human.brain-map.org/static/download). Subjects had no history of neuropsychiatric or neuropathological disorders (demographics are tabulated in Supplementary Table 12). Tissue samples were collected for microarray analysis by either manual macrodissection for large regions (cortical and subcortical structures) or by laser-based microdissection for smaller regions (subcortical and brainstem areas)[12]. An average of 466 samples (left hemisphere) were obtained from four donor brains (466 ± 72.6 samples from H0351.1009, H0351.1012, H0351.1015, and H0351.1016). The remaining two donor brains (H0351.2001 and H0351.2002) supplied 946 and 893 samples, respectively, covering both hemispheres. Cortical samples from the left hemisphere were included in the current study[13]. For each sample, RNA isolation, quantification, normalization, and quality control were performed. Microarray analysis was conducted by Beckman Coulter Genomics company (for details, see “Technical White Paper: Microarray Survey” in http://help.brain-map.org/display/humanbrain/Documentation). The normalized expression levels of 58,692 probes representing 20,737 genes were subsequently obtained. We updated gene symbols by replacing the previous and alias gene symbols by the approved symbols from the HUGO Gene Nomenclature Committee (HGNC) database (http://biomart.genenames.org/). For each donor brain, expressions of probes corresponding to the same gene symbol were averaged, resulting in an array containing 20,734 expression levels for each sample. Gene expression levels were further normalized within each sample by dividing expression values by the mean expression of the sample.

Next, tissue samples were spatially mapped to FreeSurfer cortical regions in order to obtain region-wise gene expression profiles, using an approach similar to the method proposed in a prior study[14]. First, the sample annotation data, including the Montreal Neurological Institute (MNI) coordinates and the structure type of each sample, was extracted in the dataset downloaded from AHBA website. Samples annotated outside the left hemisphere of cerebral cortex were excluded. Second, FreeSurfer software was applied to process the MNI-152 template for brain tissue segmentation and cortical mantle reconstruction[3]. The reconstructed cortical mantle was parcellated into the distinct cortical regions of the atlas used (i.e., DK-114 with 57 regions per hemisphere based on the Cammoun subdivision of the Desikan-Killiany atlas[4, 6, 7]). A finer subdivision of the Desikan-Killiany atlas containing 219 regions (111 regions on the left hemisphere) was used for a validation, as well as the cytoarchitectonic chimpanzee-human homologous BB-38 atlas (see below). Third, for each sample in the AHBA data, the nearest voxel in the MNI 152 template was searched, according to the Euclidean distance computed between MNI coordinates of the AHBA sample and all gray matter voxels in the MNI 152 template. Each sample was assigned to a cortical region based on the nearest gray matter voxel. A distance threshold of 2 mm was used to exclude inaccurate assignments of cortical regions (also 1 mm and 3 mm were examined for validation). The assignment was manually verified to ensure that no subcortical sample was included. Finally, for each donor brain, gene expression profiles of samples belonging to the same cortical region were averaged, resulting in a 6 × 57 ×

20,734 data matrix (i.e., donors × cortical regions × genes). Within each donor, gene expressions were normalized to $Z$ scores across all cortical regions per gene. The normalized gene expression profiles were averaged across 6 donor brains to obtain a group-level gene expression matrix of a size of 57 × 20,734.

#### *HAR identification using brain-related Hi-C and eQTL*

The set of HAR genes as used in the main text was obtained from the study of Doan et al.[15], as those where HARs are located within the introns, within or near (less than 1kb) 5′ and 3′ UTRs, or are the closest flanking gene that was less than 2.1mb away, (with 70% being less than 500kb away)[15]. An alternative approach to identify HAR genes can be gene mapping based on brain related chromatin interaction or eQTL. PsychENCODE provides variety of functional genomic datasets[16] (http://resource.psychencode.org/). We obtained significant eQTLs and enhancer-promoter (gene) linkages based on HiC in prefrontal cortex from http://resource.psychencode.org/. In both dataset, genes were provided by ensemble gene ID. We filtered on protein coding genes based on Ensembl v92 GRCh37. HAR regions overlapped with significant eQTLs were mapped to the genes whose expression is potentially affected by the SNPs in HAR region. Similarity enhancer regions were overlapped with HAR regions and mapped to genes whose promoter is interacting with the potential enhancer.

#### *Functional networks*

Resting-state functional networks were obtained from the Yeo 7-network atlas[17]. The Yeo atlas contains a parcellation map of 7 large-scale functional resting-state networks, including the visual, somatomotor, dorsal attention, ventral attention (also referred to as salience), limbic, frontal-parietal (FPN; also referred to as central executive), and default mode network (DMN). An annotation file of the 7 functional networks was included for the fsaverage subject in the FreeSurfer Software package (https://surfer.nmr.mgh.harvard.edu/fswiki/CorticalParcellation_Yeo2011). We assigned each cortical region in the Desikan-Killiany atlas[6, 7] (BB-38 atlas for validation purposes) to one of the 7 functional networks. For this, the surface-based annotation was translated to a 3D brain volume in volumetric space, in which each gray matter voxel was assigned a network label. Next, for each region in the Desikan-Killiany atlas, we computed the ratio of voxels that belonged to each of the 7 functional networks. Using majority vote, the label of the functional network containing the majority of voxels was then assigned to that region. The original vertex-based functional networks are shown in Supplementary Fig. 2 and the regional mapped functional network assignment is shown in Fig. 2b in the main text. For a validation analysis, DMN regions were additionally cross-referenced to the definition of the DMN by Raichle et al for a validation[18], resulting in 14 DK regions forming the DMN (Supplementary Fig. 10). A finer parcellation of the Yeo atlas in 17 networks[17] was also used for validation purposes (Supplementary Fig. 11).

***UK Biobank MRI and GWAS on DMN functional connectivity***

The UK Biobank study protocol was approved by the National Research Ethics Service Committee North West– Haydock (reference 11/NW/0382), and all procedures were conducted in accordance with the ethical principles for medical research declared in the World Medical Association Declaration of Helsinki. Access to the UK Biobank data was obtained under the UK Biobank application number 16406. All participants were recruited by invitation letters that were sent out to approximately 9.2 million individuals between 37-72 years living within 25 miles distance from one of the 22 study assessment centers and were registered with the National Health Service (NHS). Data collected included genotype data ascertained from blood samples and a wide array of phenotypic data, such as registry-based phenotypic information, extensive self-reported baseline data collected by questionnaire and brain imaging, among others.

*Neuroimaging data and processing*

The UK Biobank MRI data were acquired on a 3 Tesla Siemens Skyra VD13A SP4 scanner with a standard Siemens 32-channel RF receive head coil. Data included the acquisition of an anatomical T1-image (parameters: 3D MPRAGE, in-plane acceleration iPAT=2, pre-scan-normalize, 1 mm isotropic voxel size, defaced, 5 minutes, T1_orig_defaced). The anatomical T1-weighted image was parcellated into 114 cortical regions according a subdivision of the Desikan-Killiany atlas[6, 7] using FreeSurfer[3]. Resting-state fMRI data were acquired according to the following parameters: GE-EPI with ×8 multi-slice acceleration, 6 minutes, 490 timepoints, TR = 735 ms, TE = 39 ms, no iPAT, flip angle 52°, and fat saturation. The fMRI data was pre-processed by the UK biobank with motion correction using MCFLIRT[19], grand-mean intensity normalization of the entire 4D dataset by a single multiplicative factor, high-pass temporal filtering (Gaussian-weighted least-squares straight line fitting, with sigma = 50.0s), EPI unwarping (using acquired fieldmaps), and gradient distortion correction (GDC unwarping). The fMRI images were aligned with the T1-weighted image using FLIRT[20]. Structured artefacts in the fMRI images were removed by ICA + FIX processing (Independent Component Analysis followed by FMRIB's ICA-based X-noiseifier[21, 22]). ICA + FIX performs rigid-body head motion correction and drift removal (by high-pass temporal filtering). Based on a hand-trained classifier ICA+FIX identified bad ICA time courses and regressed these bad ICA time courses and 24 motion parameters from the data. We further processed the fMRI data by removing nuisance effects (regressing out the mean signal of white matter and ventricle, as well as 6 motion parameters) and by band-pass filtering (0.01 - 0.1 Hz). Potential effects of motion were removed by means of ‘scrubbing’, removing scan frames from the individual time-series in which significant movement was detected[23]. Functional connectivity between each pair of the 114 cortical regions was assessed by means of correlation analysis, computed as the Pearson correlation coefficient between the mean time-series of all region pairs.

*Genetic data and processing*

Genome-wide association study (GWAS) was performed based on a cohort of 6,849 participants from UK Biobank (http://www.ukbiobank.ac.uk)[24]. Details on all subsequently described genetic procedures and quality control are described previously (Savage et al.[25]) and include the following steps:

Imputed genotype data were obtained from the second release by UK Biobank (July 2017), including 92,693,895 genetic variants in 487,442 individuals[26]. In this study, we specifically used the data of 6,849 individual of which neuroimaging data (i.e., T1-weighted MRI, resting-state fMRI, and diffusion-weighted imaging) were also available at the time. Genotyping was divided over 106 batches using two custom Affymetrix genotyping platforms (UK BiLEVE Axiom array n ~= 50,000; UK Biobank Axiom array n ~= 450,000). Quality control of the genotype data was performed locally by the UK Biobank (details available at http://www.biorxiv.org/content/early/2017/07/20/166298). Genotypes were imputed using a combination of two reference panels: 1) a merged reference panel that included the UK10K haplotype panel and 2) the 1000 Genomes reference panel. In addition, the genotype data was imputed using the Haplotype Reference Consortium (HRC) reference panel[27]. If variants were imputed in both panels, the HRC imputation was retained.

In the current GWAS, only individuals of European ancestry were included, defined by projecting ancestry principal components from the 1000 Genomes reference populations[28] onto the called genotypes available in UK Biobank, and classifying individuals into their closest ancestral population according to the minimum Mahalanobis distance from the projected principal component scores[29]. We excluded subjects with a Mahalanobis distance > 6 standard deviation (SD) from their empirically assigned population. Additional filtering of individuals was based on UKB-provided information on genomic relatedness (subjects with most inferred relatives, 3rd degree or closer, were removed until no related subjects were present), discordant sex, sex aneuploidy, missing phenotype or covariate data, and withdrawn consent.

Imputed variants were converted to hard calls at a certainty threshold of 0.9, filtering by an imputation INFO threshold of < 0.9 and excluding multi-allelic SNPs, indels, SNPs without a unique rsID, and SNPs with minor allele frequency (MAF) < 0.0073 (i.e., minor allele count < 100), resulting in a total of 9,395,645 remaining SNPs for analysis. To correct for population stratification, we computed European-specific principal components based on a set of 145,432 independent ($r^2 < 0.1$) autosomal SNPs with MAF > 0.01 and INFO = 1 using FlashPCA2[30].

A Genome-wide analysis was performed on functional connectivity strength of the default-mode network (DMN-FC). DMN-FC was computed as the ratio between the sum of FC strength between regions in the DMN and the sum of functional connectivity between all other regions of the brain. GWAS was conducted in PLINK v1.9[31] using an additive linear regression model and controlling for covariates of age, sex, twenty European-based ancestry principal components, and total brain volume (computed for each individual as the sum of volume of grey and white matter provided by FreeSurfer[3]).

Details on all subsequently described genetic procedures and quality control were taken from the study of Savage and colleagues[25]: all files were checked for data integrity and accuracy. SNPs were filtered from further analysis if they met any of the following criteria: imputation quality (INFO/$R^2$) score < 0.6, Hardy–Weinberg equilibrium $p < 5 \times 10^{-6}$, minor allele frequency (MAF) corresponding to a minor allele count (MAC) < 100, and mismatch of alleles or allele frequency difference greater than 20% from the Haplotype HRC genome reference panel[27]. Indels and SNPs that were duplicated, multiallelic, monomorphic, or ambiguous (A/T or C/G with MAF > 0.4) were also excluded. Visual inspection of the distribution of the summary statistics was performed, and Manhattan plots and quantile–quantile plots were created for the cleaned summary statistics.

*Genomic locus definition.* Independently associated loci resulting from the GWAS on DMN-FC were further defined using FUMA[32]. Independent significant SNPs, which were identified by a Bonferroni-corrected genome-wide significant two-tailed *p* value ($p < 5 \times 10^{-8}$), represented signals that were independent from each other at linkage equilibrium $r^2 < 0.6$. A subset of the independent significant SNPs that showed approximate linkage equilibrium with each other at $r^2 < 0.1$ were defined as 'lead SNPs'. We then defined associated 'genomic loci' by merging any physically overlapping lead SNPs (LD blocks < 250 kb apart). Borders of the associated genomic loci were defined by identifying all SNPs in LD ($r^2 \geq 0.6$) with one of the independent significant SNPs in the locus, and the region containing all of these 'candidate SNPs' was considered to be a single independent genomic locus. LD information was calculated from UKB genotype data [25].

*Functional annotation of SNPs.* Functional annotation of identified DMN-FC SNPs was examined using FUMA[32]. We selected all candidate SNPs in the associated genomic loci having $r^2 \geq 0.6$ with one of the independent significant SNPs, a suggestive $p$ value ($p < 1 \times 10^{-5}$), and MAF > 0.0073 (i.e. MAC > 50) for annotations. Predicted functional consequences for these SNPs were obtained by matching SNPs' chromosome, base-pair position, and reference and alternate alleles to ANNOVAR databases[33], obtaining ANNOVAR categories that identify the SNP's genic position (for example, intron, exon, intergenic) and associated function.

*Gene mapping.* Genome-wide significant loci obtained by the GWAS analysis were mapped to genes using FUMA[32] according to three strategies: (1) Positional mapping projects SNPs to genes based on physical distance (within a 10-kb window) from known protein-coding genes in the human reference assembly (GRCh37/hg19); (2) eQTL mapping projects SNPs to genes with which they show a significant eQTL association (i.e., allelic variation at the SNP is associated with the expression level of that gene). eQTL mapping uses information from 45 tissue types in 3 data repositories (GTEx[34], Blood eQTL browser[35], BIOS QTL browser[36]) and is based on cis-eQTLs that can map SNPs to genes up to 1 Mb away. We used an FDR of $q < 0.05$ to define significant eQTL associations; (3) Chromatin interaction mapping was performed to map SNPs to genes when there was a 3D DNA–DNA interaction between the SNP region and a gene region. Chromatin interaction mapping can involve long-range interactions, as it does not have a distance boundary. FUMA contained Hi-C data for 14 tissue types from the study of Schmitt et al.[37]. Because chromatin interactions are often defined in a certain resolution, such as 40 kb, an interacting region can span multiple genes. If a SNP is located in a region that interacts with a region containing multiple genes, it will be mapped to each of those genes. To further prioritize candidate genes, we selected only interactionmapped genes in which one region involved in the interaction overlapped with a predicted enhancer region in any of the 111 tissue/cell types from the Roadmap Epigenomics project[38] and the other region was located in a gene promoter region (from 250 bp upstream to 500 bp downstream of the TSS and also predicted by Roadmap to be a promoter region). This reduced the number of genes mapped but increased the likelihood that those identified would have a plausible biological function. We used an FDR of $q < 1 \times 10^{-5}$ to define significant interactions, based on previous recommendations[37] modified to account for the differences in cell lines used here.

*GWAS catalog lookup.* We used FUMA GENE2FUNC[32] to identify SNPs with previously reported ($p < 5 \times 10^{-5}$) phenotypic associations in 56 published GWAS listed in the NHGRI-EBI Catalog[39] that overlapped with the genomic risk loci identified in the current GWAS analysis. The enrichment was tested using a hypergeometric test with a background set of 19,283 genomic protein-coding genes as in FUMA.

*Gene-set analysis.* MAGMA (v1.07)[40] was used to perform gene-set analysis, which is based on the linear regression model, testing for associations of HAR genes with phenotypes. Conditional gene-set analyses were performed as a follow-up using MAGMA to correct for the effect of BRAIN genes. MAGMA-based gene-set analysis expands hypergeometric enrichment testing in FUMA[32] in the sense that MAGMA weighs the contribution of genes based on the association *p*-value with the trait, whereas in Hypergeometric enrichment genes are denoted as ‘implicated’ and then tested for overlap with a gene-set.

***Pathway-sharing analysis***

To examine whether genes associated with DMN-FC were functionally related to the set of HAR genes, we used the data of curated gene sets (C2) and GO gene sets (C5) of the MSigDB collections (http://software.broadinstitute.org/gsea/msigdb/collections.jsp)[41], and obtained a total number of 19,427 genes and 7,246 pathways. A gene by gene matrix, *G*, was constructed, with the element $G(i,j)$ assigned by 1 if gene *i* and gene *j* were involved in at least one pathway at the same time, otherwise assigned by 0. We calculated the sum of elements in *G* between DMN-FC associated genes and HAR genes (i.e., summing up $G$(i,j), taking gene *i* from DMN-FC relevant genes and gene *j* from HAR genes), indicating the number of gene pairs from these two gene sets sharing the same pathway. Permutation analysis was performed by constructing a null distribution, in which the same sized set of random genes was selected to replace HAR genes and the number of gene pairs sharing the same pathway was counted (10,000 permutations). The original number of gene pairs was then compared to the null distribution to achieve a *p* value.

***Top genes differentiating genes of the default-mode network***

Two-tailed two-sample *t* tests were performed for each of 20,734 AHBA genes to examine the difference in gene expression between regions of the DMN and the rest of the brain. Genes showing the top 200 largest *t* scores were selected as the DMN genes. We alternatively examined the top 53 ($p < 0.0014$, partial Bonferroni corrected) and 469 genes ($p < 0.01$, not corrected) for validation purposes. Notably, because expression levels of genes were not independent, the partial Bonferroni correction[42, 43, 44] was used in the current study, with threshold determined based on the number of principle components (n = 36) explaining 95% of the total variance of the gene expression of all genes using principal component analysis (PCA), resulting in an adjusted $p < 0.05/36 = 0.0014$.

We calculated the ratio of genes within the top DMN genes overlapping with the 1,711 HAR genes. For statistical evaluation, we tested the enrichment of HAR genes for DMN genes using hypergeometric test. To examine whether HAR genes was specifically enriched for DMN genes, a permutation analysis was performed. In each of the 10,000 permutations, functional network labels were shuffled and top 200 genes showing the largest expression differences between the reshuffled DMN and other regions were selected. The proportion of HAR genes in these top genes was computed to generate a null distribution. The original ratio was then compared to the null-distribution to obtain the proportion of random permutations that exceeded the original ratio, and a *p* value was generated accordingly.

Gene-enrichment analysis was performed for the set of top DMN genes by means of hypergeometric test[32] to examine whether DMN genes were enriched for the predefined gene sets in three functional categories, including biological process, molecular function, and cellular component, based on the Gene Ontology (GO)[45]. For each of the predefined gene sets, a *p* value was calculated based on the number of genes present in both the predefined set and the DMN gene set. The resulting *p* values were corrected for multiple testing through FDR with $q < 0.05$.

***Gene-set analysis based on GWAS of cognitive ability, social behavior, and schizophrenia***

Association results from prior GWAS studies were used to examine the potential association of HAR and DMN genes with human cognitive ability, social behavior, and schizophrenia. SNP-based summary statistics of three GWAS studies were obtained, including 1) a recent GWAS meta-analysis on intelligence in 269,867 individuals [25] (https://ctg.cncr.nl/software/summary_statistics); 2) a GWAS of a social interaction related trait, "Frequency of friend/family visits", in 383,941 individuals in the UK Biobank (GWAS ATLAS web tool, http://atlas.ctglab.nl/traitDB/3216); 3) a GWAS in 33,426 schizophrenia patients and 54,065 healthy controls[46] as provided by the Psychiatric Genomics Consortium (http://www.med.unc.edu/pgc/). Gene annotation was performed using MAGMA[40], providing the gene-based effect size that is not directional and can be in both positive and negative direction given phenotypic variants. The gene-set analysis was performed based on the linear regression model implemented in MAGMA[40] to examine to what extent HAR and DMN genes are associated with above phenotypic variations. The conditional gene-set analysis was also used to control for the effect of general BRAIN genes.

***Cortical vulnerability in psychiatric disorders***

We examined the potential role of evolutionary genes in brain morphology related to psychiatric disorders. A cortical map of disorder involvement was derived from the BrainMap database (http://www.brainmap.org/), describing a large collation of standardised data from voxel-based morphometry (VBM) studies on cortical volume changes in a wide range of brain disorders (994 studies in total)[47, 48, 49]. Five psychiatric disorders were selected (260 studies), including autism spectrum disorder, bipolar disorder, major depressive disorder, obsessive compulsive disorder, and schizophrenia, for which 519, 547, 751, 215, and 2840 disorder-hit voxels (with MNI coordinates) were included. Meta-analyses were conducted for each disorder using the GingerALE toolbox[50, 51, 52]. Resulting brain maps of activation likelihood estimation (ALE) were registered to the MNI 152 template in the Freesurfer space and regional ALE was computed by averaging ALEs of all voxels within each cortical region of the DK-114 atlas. Regional averaged

ALE was transformed to *Z*-score and then averaged into a cross-disorder cortical involvement map describing per region the level of involvement across the five major psychiatric disorders.

### Supplementary Results

#### *HAR genes identified by brain-related Hi-C and eQTL*

The genomic HAR segments as described by Doan et al.[15] were mapped to 184 genes using the eQTL mapping and chromatin interaction mapping defined in brain tissues from the PsychENCODE database[16]. Using these 184 HAR genes, we found that the mean cortical gene expression profile was significantly correlated to the cortical expansion from chimpanzees to humans ($r(53) = 0.321$, $p = 0.017$), but the correlation did not exceed the null distribution of correlation coefficients between gene expression of random ECE genes and cortical expansion ($p = 0.184$, 10,000 permutations). Furthermore, cortical gene expression of regions in higher-order cognitive networks (i.e., DMN, FPN, and VAN) were significantly higher than regions of SMN/VN ($t(44) = 2.883$, $p = 0.010$), with the DMN showing significantly higher gene expression compared to the rest of the cortex ($t(55) = 2.157$, $p = 0.035$). These effects significantly exceeded null distributions of effects obtained by randomly selecting the same sized sets of ECE genes ($p = 0.009$ and 0.019, separately; 10,000 permutations).

We further examined a subset of genes (32 out of the 184 HAR genes) that was overlapped with the 2,979 BRAIN genes. First, cortical gene expression of these 32 HAR-BRAIN genes was significantly correlated to the cortical expansion from chimpanzees to humans ($r(53) = 0.405$, $p = 0.002$), but this correlation again did not exceed null distributions generated by either random BRAIN genes (NULL1, $p = 0.187$) or random ECE genes (NULL2, $p = 0.052$). Moreover, regions of higher-order cognitive networks showed elevated gene expression as compared to regions of the SMN/VN ($t(44) = 4.026$, $p < 0.001$), with the effect exceeding NULL2 ($p = 0.030$, 10,000 permutations), but not NULL1 ($p = 0.255$, 10,000 permutations). Examining the potential elevation of gene expression in the DMN regions compared to the rest of the cortex also showed a significant effect ($t(55) = 3.240$, $p = 0.002$), exceeding effects from both NULL1 ($p = 0.009$, 10,000 permutations) and NULL2 ($p = 0.003$, 10,000 permutations). These findings suggested that the 32 HAR-BRAIN genes identified according to brain eQTL and Hi-C might play a specific role in differentiating the DMN from other functional networks.

#### ***HAR genes identified by MPRA***

Using massively parallel reporter assays (MPRA), Doan et al.,[15] identified biallelic HAR mutations related to autism spectrum disorder and mapped these genomic loci to 238 genes according to genomic locations (here further referred to as HAR-ASD genes). In the current study, we used the original list of 238 genes in Doan et al., and did not combine the MPRA results with brain Hi-C/eQTL as it only resulted in 4 genes, which did not leave enough statistical power to pick up any effects. First, cortical gene expression profile of these HAR-ASD genes was found to be correlated to cortical expansion from chimpanzees to humans ($r(53) = 0.410$, $p = 0.002$), with the correlation coefficient significantly larger than null distribution of correlations between cortical expansion and gene expression of random ECE genes ($p = 0.011$, 10,000 permutations). Concerning the seven functional networks, higher-order cognitive networks showed enhanced gene expression of HAR-ASD genes as compared to regions of the SMN/VN ($t(44) = 3.701$, $p < 0.001$), with the DMN regions showing significantly higher gene expression comparing to the rest of the cortex ($t(55) = 2.539$, $p = 0.014$). These effects significantly exceeded effects obtained from random ECE genes (both $p < 0.001$, 10,000 permutations).

Within the 238 HAR-ASD genes, 63 genes were observed to be also described as BRAIN genes (referred to as HAR-ASD-BRAIN genes). The mean cortical gene expression of these 63 genes were significantly correlated to cortical expansion from chimpanzees to humans ($r(53) = 0.484$, $p < 0.001$), exceeding correlations from both NULL1 (i.e., BRAIN genes; $p = 0.021$, 10,000 permutations) and NULL2 (i.e., ECE genes; $p = 0.004$, 10,000 permutations). Furthermore, regions of the higher-order cognitive networks showed an elevated gene expression of HAR-ASD-BRAIN genes as compared to regions of the SMN/VN ($t(44) = 5.871$, $p < 0.001$), an effect significantly larger than effects from NULL1 ($p = 0.031$, 10,000 permutations) and NULL2 ($p = 0.002$, 10,000 permutations). The DMN regions also showed higher gene expression of HAR-ASD-BRAIN genes compared to the rest of the cortex ($t(55) = 3.234$, $p = 0.002$), again, showing a larger effect compared to effects of NULL1 ($p = 0.002$, 10,000 permutations) and NULL2 ($p = 0.004$, 10,000 permutations). These findings suggested that the subset of HAR genes associated with ASD de novo variants might be involved in the process differentiating regions of cognitive systems, in particular the DMN.

***Top genes differentiating the default-mode network***

In the main text the analysis of the top 200 DMN genes is discussed. For validation, the top 53 and 469 genes were examined, showing respectively the highest positive $t$ scores (with $p < 0.0014$, partial Bonferroni corrected, and $p < 0.01$, not corrected) between the expression level in regions of the DMN and the rest of the cortex. Out of the top 53 and 469 genes (from now on referred to as 53|469), 15|68 were observed in HAR genes (both $p < 0.001$, hypergeometric test), with 6|32 genes found in the subset of HAR-BRAIN genes (both $p < 0.001$, hypergeometric test). Permutation analysis by shuffling region labels across the seven functional networks revealed similar findings as described in the main text ($p < 0.001$, for both top 53 and 469 DMN genes; 10,000 permutations), suggesting an enrichment of HAR genes in the top DMN genes. Using the

the hypergeometric test[32], we again observed neuron-related GO annotations for the top 53|469 DMN genes, with results tabulated in Supplementary Table 13 and 14.

We further examined to what extent the top 53|469 DMN genes were associated with the genetic variants of intelligence, social behaviour, and schizophrenia. As a result, the top 53 DMN genes were found to be significantly associated with genetic variants of Frequency of friend/family visits ($ß = 0.013$, $p = 0.022$) and schizophrenia ($ß = 0.017$, $p = 0.014$); no clear significant effect for intelligence could be observed ($ß = -0.001$, $p = 0.568$). The top 469 DMN genes showed significant association with all three phenotypes (intelligence: $ß = 0.022$, $p = 0.004$; Frequency of friend/family visits: $ß = 0.015$, $p = 0.008$; schizophrenia: $ß = 0.015$, $p = 0.026$).

***Examination of the potential effects of tissue sample number***

In the AHBA dataset used in the principal analysis, a distinct number of tissue samples was obtained within each of the 114 DK regions (mean [SD] = 3.3 [2.5], ranging from 1 ~ 13). Gene expression profiles of tissue samples were averaged within brain regions across different numbers of tissue samples. Here we verified our key results of HAR-BRAIN gene expression considering the potential effects of variations in the number of tissue samples in each cortical region.

First, we regressed out the number of tissue samples from the gene expression profile using the linear regression model and found similar results using the residuals of gene expression. We found that the HAR-BRAIN gene expression was significantly correlated with cortical expansion ($r(55) = 0.491$ , $p < 0.001$; $p < 0.001$ for both NULL1 and NULL2, 10,000 permutations). Moreover, regions in higher-order cognitive networks (i.e., DMN, FPN, and VAN combined) showed higher HAR-BRAIN gene expression as compared to the SMN/VN (t(44) = 4.657, $p$ < 0.001; $p$ = 0.013 and $p$ < 0.001 for NULL1 and NULL2, respectively, 10,000 permutations), with the DMN regions showing the highest gene expression compared to the rest of the cortex (t(55) = 3.154, $p$ = 0.003; $p$ < 0.001 for both NULL1 and NULL2, 10,000 permutations).

Second, we performed 1,000 randomizations, in which a single tissue sample was randomly selected per region, and the gene expression profile of the selected sample was used to represent the profile of the corresponding DK region. Across 1,000 randomizations, we observed an averaged $t(44) = 4.401$ ($p < 0.001$) when comparing gene expressions of the 415 HAR-BRAIN genes between regions in cognitive and primary networks, as well as an averaged $t(55) = 2.436$ ($p = 0.018$) between regions of the DMN and the rest of the cortex. Furthermore, the pattern of HAR-BRAIN gene expression was correlated to the pattern of chimpanzee-to-human cortical expansion, with an averaged correlation coefficient of $r = 0.369$ ($p = 0.005$). These findings confirmed the high-level HAR-BRAIN gene expression in cognitive functional networks and its association with evolutionary cortical expansion.

#### *Alternative cortical parcellation atlases*

*BB-38 atlas for chimpanzee-human comparison.* We validated the association between cortical evolutionary expansion and HAR gene expression using the BB-38 atlas that describes homologous cortical areas across the two species. Cortical expansion of the seven resting-state functional networks were displayed in Supplementary Fig. 3a, showing the largest expansion in the FPN and the second largest in the DMN. Regions of the higher-order cognitive networks demonstrated larger expansion as compared to the SMN/VN ($t(55) = 3.632$, $p < 0.001$; Supplementary Fig. 3b). Furthermore, we found higher gene expression levels for the set of

HAR-BRAIN genes in regions of cognitive networks comparing to the SMN/VN ($t(28) = 2.572$; $p = 0.016$; Supplementary Fig. 3b). Cortical gene expression of HAR-BRAIN genes was significantly correlated with cortical evolutionary expansion ($r = 0.400$, $p = 0.014$, Supplementary Fig. 3c), suggesting consistent findings between the BB-38 atlas and DK atlas.

*DK-219 atlas*. Results were further validated using a finer subdivision of DK atlas consisting of 219 cortical regions (111 regions in left hemisphere)[6, 7]. Regions of the DMN, FPN, and VAN consistently showed larger cortical surface area expansion as compared to regions of the SMN/VN ($t(171) = 3.671$, $p < 0.001$). A similar effect was also observed when we compared regions of the DMN with the SMN/VN ($t(124) = 2.895$, $p = 0.005$). The pattern of chimpanzee-to-human cortical expansion consistently correlated with the pattern of gene expression for HAR-BRAIN genes ($r(109) = 0.390$, $p < 0.001$), again, significantly exceeding null conditions in which the similar sized set of BRAIN genes were randomly selected ($p < 0.001$ for HAR-BRAIN genes; 10,000 permutations). A significantly enhanced HAR-BRAIN gene expression remained in cognitive network regions as compared to the SMN/VN regions ($t(84) = 5.293$, $p < 0.001$), as well in the DMN regions as compared to the rest of the cortex ($t(109) = 2.247$, $p = 0.027$), with effects remaining significant in permutation testing conditional to BRAIN genes ($p = 0.016$ and $p < 0.001$, separately, 10,000 permutations).

***Alternative functional network divisions***

*Yeo-2011 17 functional networks*. We validated our results using a finer functional network division that consists of 17 functional networks[17]. Four out of 17 functional networks (labeled as 11, 15, 16, and 17 in [17]) were identified as components of the DMN, as shown in Supplementary

Fig. 11. A group of these DMN components and networks labeled as 7, 8, 12, and 13 was noted as higher-order cognitive networks, meanwhile networks labeled as 1, 2, 3, and 4 were defined as low-order primary networks. Using this functional network division, a significantly larger cortical expansion in regions of higher-order cognitive networks as compared to primary networks was confirmed ($t(82) = 2.185$, $p = 0.032$). Regions in cognitive networks consistently exhibited higher gene expression for HAR-BRAIN genes ($t(40) = 5.182$, $p < 0.001$), as compared to regions of the SMN/VN. Comparing the DMN regions to the rest of the cortex showed similar results ($t(55) = 2.306$, $p = 0.025$). These effects remained to be significant in permutation testing conditional to BRAIN genes ($p = 0.009$ and $p = 0.003$, separately, 10,000 permutations).

*Customized default-mode network.* We additionally validated our findings by using a refined DMN division according to Raichle[18]. We manually identified 14 cortical regions in DK atlas as default-mode network regions, including regions located in precuneus, middle temporal lobe, inferior parietal lobe, and medial frontal lobe (Supplementary Fig. 10). Under this division, we consistently observed significantly enhanced HAR-BRAIN gene expression in the DMN regions as compared to the rest of the cortex ($t(55) = 2.476$, $p = 0.016$). Permutation analysis by randomly selecting the similar sized set of BRAIN genes also showed consistent results ($p$ = 0.003, 10,000 permutations).

#### *Directly mapping AHBA tissue samples to functional networks*

We validated our findings by directly mapping the ~400 tissue samples of each donor in AHBA dataset to the functional networks, without the intermediate step of mapping both sets to the DK-114 atlas. Gene expression profiles of tissue samples within the same functional network were averaged, resulting in a 7 × 20,734 matrix representing the mean gene expression for each gene per functional network. A 7 × 1,711 sub-matrix of gene expressions of HAR genes was further selected. We then examined whether HAR genes showed the highest expression level in the default-mode network by performing paired-sample two-sided *t* tests across HAR genes between the DMN and each of other 6 functional networks. This analysis showed the highest HAR gene expressions in the DMN, as compared with the visual ($t(1{,}710) = 2.517$, $p = 0.012$), somatomotor ($t(1{,}710) = 10.369$, $p < 0.001$), dorsal-attention ($t(1{,}710) = 2.294$, $p = 0.021$), ventral-attention ($t(1{,}710) = 11.267$, $p < 0.001$), limbic ($t(1{,}710) = 2.392$, $p = 0.017$, and frontal-parietal networks ($t(1{,}710) = 7.844$, $p < 0.001$). Similar findings were observed when we considered the subset of 415 HAR-BRAIN genes ($t(414) = 3.682$, $p < 0.001$, visual; $t(414) = 10.323$, $p < 0.001$, somatomotor; $t(414) = 2.394$, $p = 0.017$, dorsal-attention; $t(414) = 7.463$, $p < 0.001$, ventral-attention; $t(414) = 5.873$, $p < 0.001$, frontal-parietal, except for the limbic network, $t(414) = -0.419$, $p = 0.676$).

Given the 7 × 415 (function networks × HAR-BRAIN genes) expression matrix, we next averaged gene expression levels across the default-node, frontal-parietal, and ventral-attention networks, resulting in a 1 × 415 vector of expression levels of HAR-BRAIN genes in higher-order cognitive functional networks. Similarly, gene expressions were averaged across primary visual and somatomotor networks. Paired-sample *t*-test was performed between the two resulting gene expression profiles and showed that HAR-BRAIN gene expression was significantly higher in cognitive networks as compared with regions of the SMN/VN ($t(414) = 5.345$, $p < 0.001$)). Also, comparing the gene expression profile of the DMN with the mean profile of other 6 functional networks showed significantly higher default-mode HAR-BRAIN gene expression ($t(414) = 8.922$, $p < 0.001$). We further performed a permutation test by selecting a same-sized set of random BRAIN genes from the original 7 × 20,734 expression matrix and recomputing the paired-sample *t* test between the mean expression profiles of higher-order cognitive and primary networks, as well as between the default-mode network and others, for 10,000 times. The original *t* scores significantly exceeded 10,000 permutations for both conditions ($p = 0.003$ for the comparison between cognitive and primary networks; $p < 0.001$ for the comparison between the DMN and others). Taking together, this validation analysis confirmed that cognitive functional networks, in particular the DMN showed the highest level of HAR/HAR-BRAIN gene expression.

***Alternative parameters in processing***

*Distance threshold for tissue sample inclusion.* In the main analysis, a distance threshold of 2 mm was used to map tissue samples in the AHBA data to the gray matter. We alternatively used thresholds of 3 mm and 1 mm to validate our findings. For both thresholds, regions in higher-order DMN, FPN, and VAN consistently showed higher HAR-BRAIN gene expression as compared to the SMN/VN regions ($t(44) = 3.911$, $p < 0.001$, and $t(44) = 5.522$, $p < 0.001$, for 1mm and 3mm, respectively). Comparing the DMN regions to the rest of the cortex showed similar results ($t(44) = 2.897$, $p = 0.054$, and $t(55) = 2.790$, $p = 0.072$, for 1mm and 3mm, respectively). Permutation analyses by selecting similar sized sets of random BRAIN genes consistently showed significance ($p = 0.014$ and 0.009, for threshold 1mm and 3 mm, separately, in the comparison between cognitive and the SMN/VN regions; $p < 0.001$ for both thresholds in the comparison between default-mode network regions and others; 10,000 permutations). Furthermore, the cortical expression pattern of HAR-BRAIN genes was also significantly correlated with chimpanzee-to-human cortical expansion ($r(55) = 0.334$ and 0.363, $p = 0.011$ and

0.006, for 1mm and 3 mm, respectively), with consistent results observed when performing permutation testing ($p < 0.001$ for both thresholds, 10,000 permutations). In addition, the cortical expression pattern of HAR-BRAIN genes were consistently associated with the pattern of cortical vulnerability to psychiatric disorders for both settings ($r(55) = 0.300$, $p = 0.026$, and $r(55) = 0.319$, $p = 0.018$, for 1 mm and 3mm, respectively). Effects were significantly larger than seen in the null condition of randomly selected BRAIN genes ($p = 0.006$ and $p = 0.030$, for 1mm and 3 mm, respectively; 10,000 permutations).

*FDR correction in BRAIN genes selection.* In addition to setting FDR $q < 0.05$ for selecting GTEx BRAIN genes in the main text, we alternatively applied FDR $q < 0.01$ and $q < 0.001$ to identify BRAIN genes from the GTEx dataset. At these two thresholds, respectively 2,544 and 2,102 genes were identified as BRAIN genes, from which 368 and 305 genes could be denoted as HAR-BRAIN genes. For both thresholds, higher-order cognitive network regions remained to show higher HAR-BRAIN gene expression as compared to the SMN/VN regions ($t(44) = 5.155$, $p < 0.001$ and $t(44) = 5.317$, $p < 0.001$, for FDR $q < 0.01$ and $q < 0.001$, respectively), with similar effects found when we compared regions of the DMN to the rest of the cortex ($t(55) = 3.269$, $p = 0.002$ and $t(55) = 3.330$, $p = 0.002$, for FDR $q < 0.01$ and $q < 0.001$, respectively). These effects remained significant in permutation testing in which random BRAIN genes were selected ($p = 0.008$ and 0.013, for FDR $q < 0.01$ and 0.001, separately, in the comparison between cognitive and primary networks; $p < 0.001$, for both thresholds, in the comparison between the DMN and others). Correlating the pattern of HAR-BRAIN gene expression to the pattern of cortical expansion consistently showed significant correlations ($r(55) = 0.392$, $p = 0.003$ and $r(55) = 0.382$, $p = 0.003$, for FDR q < 0.01 and 0.001, separately), with both effects significant in permutation testing conditional to BRAIN genes ($p$ = 0.001 for both thresholds, 10,000 permutations). Furthermore, cortical HAR-BRAIN gene expression was significantly associated with the cortical pattern of disorder vulnerability ($r(55) = 0.354$, $p = 0.008$ and $r(55) = 0.327$, $p = 0.015$ for FDR q < 0.01 and 0.001, respectively), effects again exceeding effects of randomly selected BRAIN genes ($p$ = 0.016 and 0.049 for $q < 0.01$ and 0.001, respectively).

***HAR-BRAIN gene expression in subcortical regions***

In the main text, we demonstrated differentiated gene expression of HAR-BRAIN genes in cortical regions of higher-order cognitive networks. Here we additionally examined whether HAR-BRAIN genes showed differentiated gene expression in cortical regions as compared to subcortical regions. To this end, we mapped subcortical AHBA tissue samples to seven subcortical regions delineated in the DK atlas, including thalamus, caudate, putamen, pallidum, hippocampus, amygdala, and nucleus accumbens. We noticed that subcortical regions showed a significantly lower HAR-BRAIN gene expression as compared to cortical regions ($t(62) = 3.427$, $p = 0.001$). This effect exceeded null distribution of effects obtained by random ECE genes (NULL1, $p < 0.001$, 10,000 permutations), but did not exceed null distribution of effects obtained by random BRAIN genes (NULL2, $p = 0.712$, 10,000 permutations). This suggested that HAR-BRAIN genes contributed to differentiating cortical regions from subcortical regions to a similar extent as genes generally involved in brain processes.

***The potential age and sex effect on expansion***

*Age*. Our study used MRI data of 29 adult chimpanzees with a mean (SD) age of 30.2 (12.6) and 30 adult humans with a mean (SD) age of 30.2 (3.1). Here we examined the potential age effect on our main cortical expansion results. We first normalized the age within each species, separately, by dividing the age of humans by 80 (an approximately mean age of human lifespan) and dividing the age of chimpanzees by 40 (an approximately mean age of captive chimpanzees). Then we regressed out the age from the surface area of each region using the linear regression model and computed cortical expansion between the two species. We found that regions of higher-order cognitive networks remained to be significantly higher than the SMN/VN regions ($t(86) = 2.667$, $p = 0.009$; FDR corrected). The FPN still showed the highest expansion (FPN versus the rest of the cortex: $t(108) = 3.092$, $p = 0.003$; FDR corrected), with the DMN ranked in the second place (DMN versus the rest of the cortex: $t(108) = 2.105$, $p = 0.038$; not corrected). Moreover, the cortical expansion significantly correlated with HAR-BRAIN gene expression ($r(53) = 0.440$, $p < 0.001$), with the correlation coefficient exceeding both NULL1 ($p = 0.012$) and NULL2 ($p = 0.002$). These findings suggest that the age effect did not drive our cortical expansion results.

*Sex*. We exclusively included MRI data of female chimpanzee and female human subjects, due to the practical reason for greater availability of female chimpanzees. Female human subjects were randomly selected from the HCP database to match the chimpanzee sample. Although it was not within the scope of the current study, it is possible that there are potential sex effects in the pattern of cortical expansion between humans and chimpanzees. Future work including sufficiently large chimpanzee samples of both male and female subjects may help to elucidate potential sex differences in cross-species cortical expansion.

#### *Permutation testing of cortical expansion*

To identify cortical regions with large chimpanzee-to-human expansion, permutation testing was performed by comparing the observed cortical expansion from chimpanzees to humans to null distributions of expansions computed after randomly shuffling the brain regions in the two species. This analysis showed significantly larger expansion in bilateral rostral middle frontal lobe, orbital inferior frontal gyrus, and right inferior/superior parietal lobe, anterior cingulate gyrus, and triangular part of inferior frontal gyrus (all uncorrected $p < 0.001$; FDR correction, $q < 0.05$; 10,000 permutations; Supplementary Fig. 4).

### Supplementary Tables

#### Supplementary Table 2. Cortical expansion in each functional network

| functional network | normalized cortical expansion | | *t* score | effect size (cohen's *d*) | adjusted *p* values |
|---|---|---|---|---|---|
| | mean | std | | | |
| VN | 0.0007 | 0.2192 | -0.6858 | -0.1724 | 0.6919 |
| SMN | -0.1000 | 0.1500 | -2.9843 | -0.7500 | 0.0122 |
| DAN | 0.0219 | 0.1803 | -0.1469 | -0.0510 | 0.8835 |
| VAN | -0.0244 | 0.1374 | -1.1175 | -0.3013 | 0.4658 |
| LN | 0.0209 | 0.2306 | -0.2150 | -0.0596 | 0.8835 |
| FPN | 0.2938 | 0.2104 | 3.4144 | 1.3321 | 0.0062 |
| DMN | 0.1167 | 0.2314 | 2.4604 | 0.5291 | 0.0359 |

#### Supplementary Table 4. HAR gene expression in each functional network

| functional network | gene expression | | *t* score | effect size (cohen's *d*) | adjusted *p* values |
|---|---|---|---|---|---|
| | mean | std | | | |
| VN | 0.0027 | 0.0621 | 0.2403 | 0.0916 | 0.8110 |
| SMN | -0.0765 | 0.0775 | -3.7018 | -1.2891 | 0.0040 |
| DAN | -0.0174 | 0.0925 | -0.3800 | -0.1970 | 0.8110 |
| VAN | 0.0051 | 0.0737 | 0.3114 | 0.1257 | 0.8110 |
| LN | 0.0165 | 0.0720 | 0.7330 | 0.2958 | 0.8110 |
| FPN | -0.0186 | 0.0609 | -0.3532 | -0.2095 | 0.8110 |
| DMN | 0.0292 | 0.0639 | 2.2737 | 0.6479 | 0.0718 |

#### Supplementary Table 5. HAR-BRAIN gene expression in each functional network

| functional network | gene expression | | *t* score | effect size (cohen's *d*) | adjusted *p* values |
|---|---|---|---|---|---|
| | mean | std | | | |
| VN | -0.0600 | 0.1145 | -1.0632 | -0.4054 | 0.4385 |
| SMN | -0.1843 | 0.1257 | -4.9206 | -1.7136 | 0.0000 |
| DAN | -0.0115 | 0.1654 | -0.0417 | -0.0216 | 0.9669 |
| VAN | 0.0024 | 0.1371 | 0.2064 | 0.0833 | 0.9419 |
| LN | 0.0596 | 0.1551 | 1.3078 | 0.5278 | 0.3535 |
| FPN | 0.0123 | 0.0584 | 0.2483 | 0.1473 | 0.9419 |
| DMN | 0.0785 | 0.0923 | 3.2673 | 0.9310 | 0.0042 |

#### Supplementary Table 7. Enrichment of DMN-FC associated genes in GWAS catalog reported gene sets

| Gene set | set size | candidates contained | adjusted *p* |
|---|---|---|---|
| Bipolar disorder | 169 | 12 | 7.35e-9 |
| Loneliness (multivariate analysis) | 23 | 4 | 2.03e-6 |

| | | | |
|---|---|---|---|
| Autism spectrum disorder or schizophrenia | 612 | 17 | 1.93e-5 |
| Iron status biomarkers (total iron binding capacity) | 36 | 4 | 2.06e-5 |
| Type 2 diabetes and other traits | 7 | 2 | 2.77e-5 |
| Venous thromboembolism | 20 | 3 | 3.24e-5 |
| Endometriosis | 41 | 4 | 3.93e-5 |
| Depressive symptoms (SSRI exposure interaction) | 10 | 2 | 9.31e-5 |
| Body mass index (SNP x SNP interaction) | 10 | 2 | 9.31e-5 |
| Cannabis dependence symptom count | 11 | 2 | 1.27e-4 |
| Tonometry | 11 | 2 | 1.27e-4 |
| D-dimer levels | 14 | 2 | 2.75e-4 |
| Systolic blood pressure response to hydrochlorothiazide in hypertension | 16 | 2 | 4.17e-4 |
| Immune response to measles-mumps-rubella vaccine | 17 | 2 | 5.02e-4 |
| Restless legs syndrome | 41 | 3 | 5.80e-4 |
| Relative hand skill in reading disability | 20 | 2 | 8.25e-4 |
| Migraine without aura | 20 | 2 | 8.25e-4 |
| Iron status biomarkers | 21 | 2 | 9.56e-4 |
| Hemostatic factors and hematological phenotypes | 21 | 2 | 9.56e-4 |
| Alzheimer's disease biomarkers | 21 | 2 | 9.56e-4 |
| Intelligence | 48 | 3 | 1.06e-3 |
| Blood osmolality (transformed sodium) | 22 | 2 | 1.10e-3 |
| Alanine aminotransferase (ALT) levels after remission induction therapy in actute lymphoblastic leukemia (ALL) | 51 | 3 | 1.33e-3 |
| Uric acid levels | 25 | 2 | 1.61e-3 |
| Age at voice drop | 25 | 2 | 1.61e-3 |

**Supplementary Table 8. Enrichment of HAR, HAR-BRAIN, BRAIN, and DMN genes in GWAS results**

| Gene set | *N* | *b* | *ß* | SE | adjusted *p* |
|---|---|---|---|---|---|
| *DMN-FC* | | | | | |
| HAR | 1509 | 0.059 | 0.016 | 0.026 | 0.020 |
| HAR-BRAIN | 377 | 0.117 | 0.016 | 0.051 | 0.020 |
| BRAIN | 2625 | 0.004 | 0.001 | 0.018 | 0.441 |
| DMN (200) | 176 | 0.081 | 0.008 | 0.063 | 0.116 |

| | | | | | |
|---|---|---|---|---|---|
| DMN-HAR-BRAIN (37) | 36 | -0.016 | -0.001 | 0.151 | 0.543 |
| *IQ* | | | | | |
| HAR | 1509 | 0.219 | 0.058 | 0.036 | < 0.001 |
| HAR-BRAIN | 377 | 0.553 | 0.076 | 0.069 | < 0.001 |
| BRAIN | 2624 | 0.173 | 0.059 | 0.024 | < 0.001 |
| DMN (200) | 176 | 0.122 | 0.011 | 0.086 | 0.098 |
| DMN-HAR-BRAIN (37) | 36 | 0.506 | 0.022 | 0.216 | 0.020 |
| *Frequency of friend/family visits* | | | | | |
| HAR | 1508 | 0.139 | 0.037 | 0.027 | < 0.001 |
| HAR-BRAIN | 377 | 0.280 | 0.039 | 0.054 | < 0.001 |
| BRAIN | 2623 | 0.052 | 0.018 | 0.018 | 0.004 |
| DMN (200) | 176 | 0.126 | 0.012 | 0.063 | 0.034 |
| DMN-HAR-BRAIN (37) | 36 | 0.320 | 0.014 | 0.159 | 0.034 |
| *Schizophrenia* | | | | | |
| HAR | 1508 | 0.075 | 0.020 | 0.032 | 0.020 |
| HAR-BRAIN | 376 | 0.305 | 0.042 | 0.063 | < 0.001 |
| BRAIN | 2622 | 0.147 | 0.051 | 0.022 | < 0.001 |
| DMN (200) | 176 | 0.124 | 0.012 | 0.077 | 0.070 |
| DMN-HAR-BRAIN (37) | 36 | 0.124 | 0.005 | 0.198 | 0.295 |

*N*: Number of genes; *b*: regression coefficient; *ß*: standardized regression coefficient; SE: standard error; *p* values adjusted for FDR.

**Supplementary Table 10. Over-representation of the top 200 DMN genes**

| Gene ontology term | *N* | *n* | adjusted *p* | gene symbol |
|---|---|---|---|---|
| *Cellular Components* | | | | |
| Dendrite | 451 | 14 | 2.60e-5 | FAS, NELL2, ITPKA, KCNJ12, SLC8A2, CHL1, GLRA3, KCNN2, PNOC, KCNB2, CALB1, NOV, NTRK2, SLC9A6 |
| Somatodendritic Compartment | 649 | 17 | 4.49e-5 | KNCN, FAS, NELL2, ITPKA, KCNJ12, SLC8A2, CHL1, GLRA3, KCNN2, VIP, PNOC, CRH, KCNB2, CALB1, NOV, NTRK2, SLC9A6 |

| | | | | |
|---|---|---|---|---|
| Cell Body | 493 | 14 | 7.15e-5 | KNCN, FAS, NELL2, KCNJ12, SLC8A2, GLRA3, KCNN2, VIP, PNOC, CRH, KCNB2, CALB1, NOV, NTRK2 |
| Perikaryon | 108 | 6 | 7.45e-5 | NELL2, SLC8A2, GLRA3, CRH, KCNB2, NTRK2 |
| Rnai Effector Complex | 11 | 2 | 1.29e-4 | DCP2, SND1 |
| Synapse Part | 607 | 15 | 2.15e-4 | SVOP, ITPKA, BAIAP3, NETO2, CBLN1, CDH8, ADCYAP1, SLC8A2, SEMA4F, LRRTM4, GLRA3, KCNN2, STX1A, CALB1, NTRK2 |
| Neuron Projection Terminus | 129 | 6 | 2.26e-4 | CDH8, ADCYAP1, PNOC, CALB1, NTRK2, SLC9A6 |
| Synapse | 751 | 17 | 2.78e-4 | OLFM3, SVOP, ITPKA, BAIAP3, NETO2, CBLN1, CDH8, ADCYAP1, SLC8A2, SEMA4F, LRRTM4, GLRA3, KCNN2, STX1A, CALB1, NTRK2, SLC9A6 |
| Cullin Ring Ubiquitin Ligase Complex | 148 | 6 | 5.20e-4 | FBXO6, KLHL12, CDC16, DCAF11, DCAF4, FBXL2 |
| T Tubule | 45 | 3 | 8.47e-4 | CACNB2, KCNJ12, KCNN2 |
| *Molecular Functions* | | | | |
| Neuropeptide Hormone Activity | 28 | 4 | 5.85e-6 | ADCYAP1, VIP, PNOC, CRH |
| Calcium Activated Potassium Channel Activity | 17 | 3 | 1.66e-5 | KCNT2, PKD2L1, KCNN2 |
| Calcium Activated Cation Channel Activity | 28 | 3 | 1.32e-4 | KCNT2, PKD2L1, KCNN2 |

*N*: Gene set size; *n*: number of contained candidates

**Supplementary Table 11. Over-representation of HAR genes**

| Gene ontology term | *N* | *n* | adjusted *p* |
|---|---|---|---|
| Neuron Part | 1261 | 525 | 1.73e-138 |
| Synapse | 751 | 378 | 2.49e-130 |
| Synapse Part | 607 | 325 | 1.52e-121 |
| Neuron Projection | 940 | 406 | 1.02e-111 |
| Postsynapse | 376 | 202 | 1.50e-75 |
| Cell Projection | 1778 | 528 | 1.35e-72 |

| | | | |
|---|---|---|---|
| Somatodendritic Compartment | 649 | 274 | 7.64e-72 |
| Axon | 417 | 210 | 9.12e-72 |
| Synaptic Membrane | 259 | 158 | 3.72e-70 |
| Presynapse | 282 | 157 | 7.77e-62 |
| Dendrite | 451 | 203 | 7.81e-59 |
| Cell Projection Part | 944 | 320 | 6.80e-57 |
| Postsynaptic Membrane | 203 | 124 | 1.71e-55 |
| Axon Part | 218 | 128 | 2.24e-54 |
| Excitatory Synapse | 196 | 117 | 6.16e-51 |
| Cell Body | 493 | 195 | 5.88e-46 |
| Transporter Complex | 318 | 147 | 1.17e-44 |
| Plasma Membrane Region | 926 | 272 | 1.91e-35 |
| Membrane Region | 1129 | 310 | 3.17e-34 |
| Cell Junction | 1146 | 311 | 2.65e-33 |

*N*: Gene set size; *n*: number of contained candidates;

Note: only top 20 cellular components tabulated

**Supplementary Table 12. Demographics of human donors included in the AHBA**

| Donor ID | Number of samples | Gender | Age (years) | Race/ethnicity |
|---|---|---|---|---|
| H0351.2001 | 946 | Male | 24 | African American |
| H0351.2002 | 893 | Male | 39 | African American |
| H0351.1009 | 363 | Male | 57 | Caucasian |
| H0351.1012 | 529 | Male | 31 | Caucasian |
| H0351.1015 | 470 | Female | 49 | Hispanic |
| H0351.1016 | 501 | Male | 55 | Caucasian |

**Supplementary Table 13. Over-representation of the top 53 DMN genes**

| Gene ontology term | *N* | *n* | adjusted *p* |
|---|---|---|---|

*Biological Processes*

| | | | |
|---|---|---|---|
| Negative Regulation of Leukocyte Chemotaxis | 13 | 2 | 3.12e-6 |
| Regulation of Monocyte Chemotaxis | 20 | 2 | 1.23e-5 |
| Regulation of System Process | 506 | 7 | 1.65e-5 |
| Adherens Junction Organization | 70 | 3 | 1.94e-5 |
| Adenylate Cyclase Activating G Protein Coupled Receptor Signaling Pathway | 72 | 3 | 2.17e-5 |
| Positive Regulation of Camp Metabolic Process | 89 | 3 | 5.00e-5 |
| Negative Regulation of Leukocyte Migration | 32 | 2 | 5.25e-5 |
| Negative Regulation of Potassium Ion Transport | 32 | 2 | 5.25e-5 |
| Positive Regulation of Vasodilation | 32 | 2 | 5.25e-5 |
| Regulation of Sensory Perception | 36 | 2 | 7.52e-5 |
| Negative Regulation of Inflammatory Response | 99 | 3 | 7.59e-5 |
| Negative Regulation of Transcription Factor Import Into Nucleus | 39 | 2 | 9.57e-5 |
| Positive Regulation of Cyclic Nucleotide Metabolic Process | 109 | 3 | 1.10e-4 |
| Positive Regulation of Adenylate Cyclase Activity | 48 | 2 | 1.79e-4 |
| Regulation of Vasodilation | 48 | 2 | 1.79e-4 |
| Negative Regulation of Chemotaxis | 50 | 2 | 2.02e-4 |
| Regulation of Camp Metabolic Process | 129 | 3 | 2.11e-4 |
| Positive Regulation of Nucleotide Metabolic Process | 132 | 3 | 2.31e-4 |
| Regulation of Myotube Differentiation | 55 | 2 | 2.68e-4 |
| Neuron Neuron Synaptic Transmission | 56 | 2 | 2.83e-4 |
| Negative Regulation of Defense Response | 143 | 3 | 3.13e-4 |
| Adenylate Cyclase Modulating G Protein Coupled Receptor Signaling Pathway | 144 | 3 | 3.21e-4 |
| Positive Regulation of Lyase Activity | 61 | 2 | 3.64e-4 |
| Homophilic Cell Adhesion Via Plasma Membrane Adhesion Molecules | 149 | 3 | 3.66e-4 |
| Negative Regulation of Response To External Stimulus | 272 | 4 | 3.73e-4 |
| Cell Cell Adhesion | 600 | 6 | 3.85e-4 |
| Negative Regulation of Response To Wounding | 155 | 3 | 4.25e-4 |

| | | | |
|---|---|---|---|
| Regulation of Cyclic Nucleotide Metabolic Process | 155 | 3 | 4.25e-4 |
| Regulation of Neurological System Process | 68 | 2 | 5.01e-4 |
| Regulation of Blood Circulation | 295 | 4 | 5.40e-4 |
| Regulation of Adenylate Cyclase Activity | 70 | 2 | 5.46e-4 |
| Single Organism Cell Adhesion | 455 | 5 | 5.52e-4 |
| Negative Regulation of Transport | 455 | 5 | 5.52e-4 |
| Negative Regulation of Nucleocytoplasmic Transport | 71 | 2 | 5.69e-4 |
| Smoothened Signaling Pathway | 72 | 2 | 5.93e-4 |
| G Protein Coupled Receptor Signaling Pathway Coupled to Cyclic Nucleotide Second Messenger | 171 | 3 | 6.15e-4 |
| Actin Mediated Cell Contraction | 73 | 2 | 6.17e-4 |
| Negative Regulation of Hormone Secretion | 74 | 2 | 6.42e-4 |
| Cell Junction Organization | 183 | 3 | 7.93e-4 |
| Cardiac Conduction | 82 | 2 | 8.66e-4 |
| Regulation of Potassium Ion Transport | 83 | 2 | 8.97e-4 |
| *Cellular Components* | | | |
| Transporter Complex | 318 | 5 | 8.00e-5 |
| T Tubule | 45 | 2 | 1.47e-4 |
| Ionotropic Glutamate Receptor Complex | 47 | 2 | 1.68e-4 |
| Calcium Channel Complex | 62 | 2 | 3.82e-4 |
| *Molecular functions* | | | |
| Neuropeptide Hormone Activity | 28 | 2 | 3.49e-5 |

N: Gene set size; n: number of contained candidates;

Gene symbol: contained gene candidates

**Supplementary Table 14. Over-representation of the top 469 DMN genes**

| Gene ontology term | *N* | *n* | adjusted *p* |
|---|---|---|---|
| *Cellular Components* | | | |
| Synapse Part | 607 | 31 | 2.93e-6 |
| Perikaryon | 108 | 11 | 3.48e-6 |
| Somatodendritic Compartment | 649 | 32 | 4.42e-6 |

| | | | |
|---|---|---|---|
| Dendrite | 451 | 25 | 5.32e-6 |
| Neuron Projection | 940 | 40 | 1.32e-5 |
| Synapse | 751 | 34 | 1.49e-5 |
| Axon Part | 218 | 15 | 2.02e-5 |
| Neuron Projection Terminus | 129 | 11 | 2.18e-5 |
| Postsynapse | 376 | 21 | 2.25e-5 |
| Terminal Bouton | 64 | 7 | 6.43e-5 |
| Cell Body | 493 | 24 | 6.78e-5 |
| Neuron Part | 1261 | 47 | 7.80e-5 |
| Axon | 417 | 21 | 1.05e-4 |
| Neuron Spine | 121 | 9 | 2.79e-4 |
| Cullin Ring Ubiquitin Ligase Complex | 148 | 10 | 3.54e-4 |
| Postsynaptic Membrane | 203 | 12 | 4.55e-4 |
| Ubiquitin Ligase Complex | 260 | 14 | 5.18e-4 |
| Cell Projection Part | 944 | 35 | 6.12e-4 |
| Myosin Filament | 22 | 3 | 1.12e-3 |
| Nuclear Replication Fork | 39 | 4 | 1.39e-3 |
| Rnai Effector Complex | 11 | 2 | 1.42e-3 |
| Filopodium Tip | 11 | 2 | 1.42e-3 |
| Cul4 Ring E3 Ubiquitin Ligase Complex | 25 | 3 | 1.84e-3 |
| *Molecular functions* | | | |
| Ubiquitin Like Protein Ligase Activity | 197 | 14 | 2.38e-5 |

*N*: Gene set size; *n*: number of contained candidates;

**Supplementary Figures**

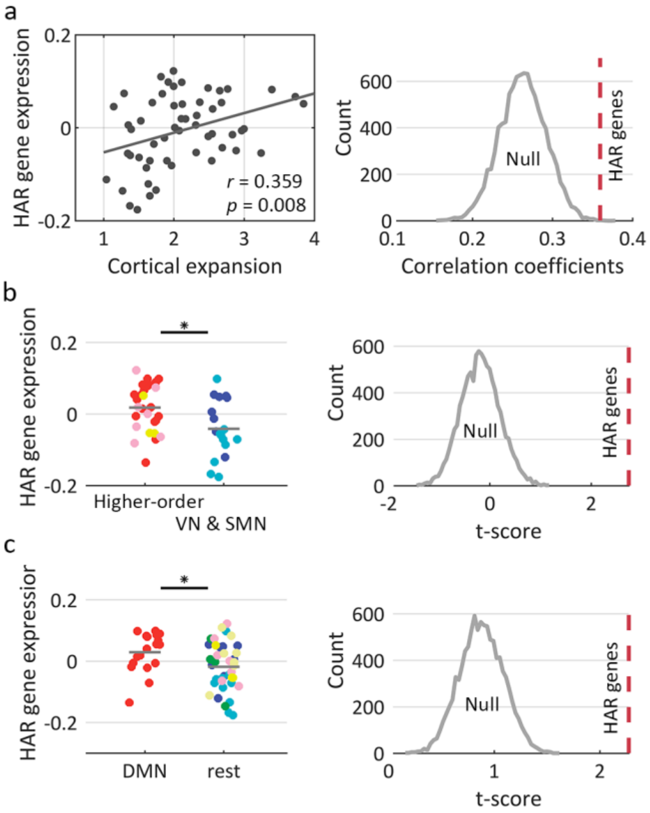


**Supplementary Figure 1.** *HAR gene expression*. (a) Cortical HAR gene expression is significantly correlated with the chimpanzee-human cortical expansion (left), with the effect significantly exceeding the null condition generated by ECE genes (right). (b) Cortical HAR gene expression in regions of higher-order networks is significantly higher than regions of the SMN/VN (left), with the effect above the null condition generated by ECE genes (right). (c) Cortical HAR gene expression in regions of DMN is significantly higher than the rest of the cortex (left), also, with the effect significantly exceeding the null condition generated by ECE genes (right). Colors indicate functional networks, as in Fig. 2 in the main text.

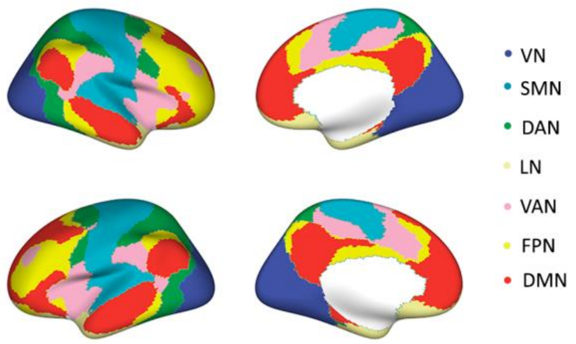


**Supplementary Figure 2.** *Brain plot of the 7 resting-state network parcellation.* VN: visual network, SMN: somatomotor network, DAN: dorsal-attention network, LN: limbic network, VAN: ventral-attention network, FPN: frontal-parietal network, DMN: default mode network.

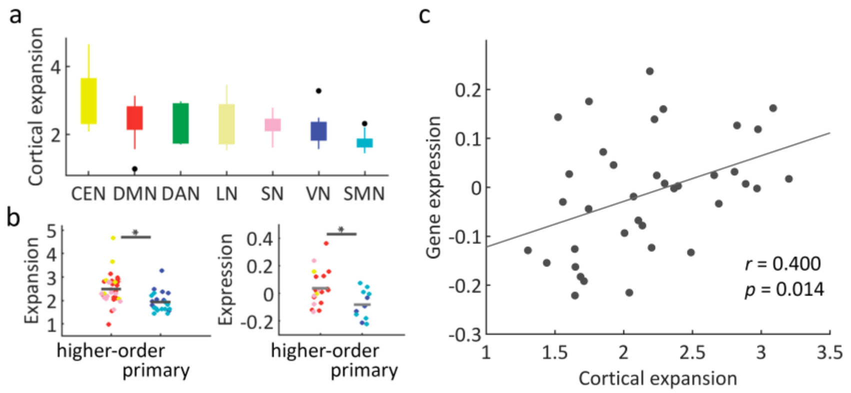


**Supplementary Figure 3.** *Evolutionary cortical expansion and HAR-BRAIN gene expression in BB-38 chimpanzee-human atlas*. (a) Levels of the cortical expansion per functional network in descending order. VN: visual network, SMN: somatomotor network, DAN: dorsal attention network, LN: limbic network, VAN: ventral-attention network, FPN: frontal parietal network, DMN: default mode network. (b) Regions of higher-order cognitive networks (DMN, FPN, VAN), present significant larger expansion (left) and higher HAR-BRAIN expression level (right), as compared to regions in primary visual and somatomotor networks (VN, SMN). (c) Cortical expansion is significantly correlated with cortical HAR-BRAIN gene expression ($r$ = 0.400, $p$ = 0.014).

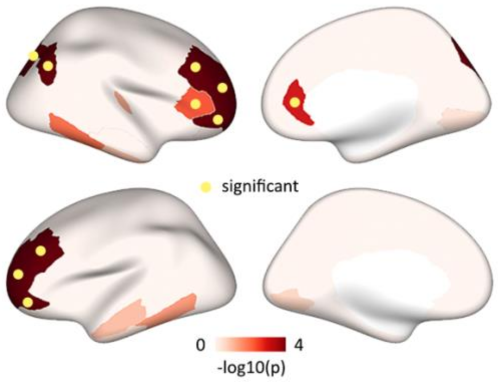


**Supplementary Figure 4.** Brain maps of regions showing significant chimpanzee-to-human expansion in permutation testing based on region shuffling.

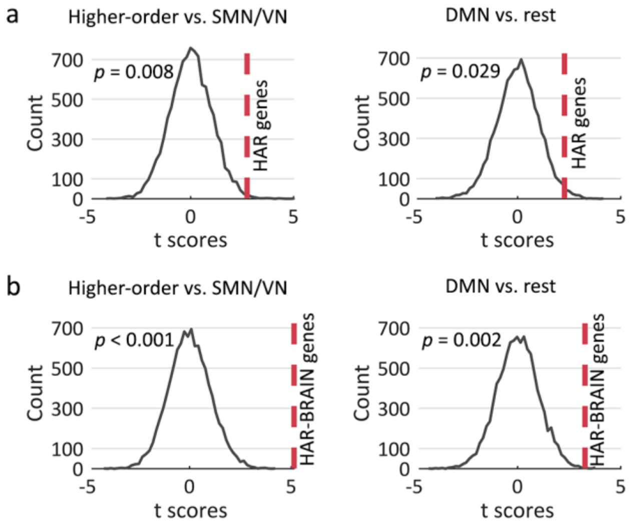


**Supplementary Figure 5**. Permutation testing by randomly shuffling cortical regions. (a) HAR gene expression between higher-order networks and the SMN/VN (left panel) and between the DMN and the rest of the cortex (right panel). (b) HAR-BRAIN gene expression between higher-order networks and the SMN/VN (left panel) and between the DMN and the rest of the cortex (right panel).

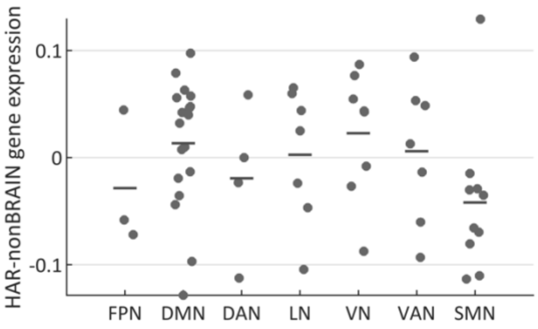


**Supplementary Figure 6**. HAR-nonBRAIN gene expression within each of the seven functional networks, ranked in descending order according to their mean expansion per network (as in Fig. 2c).

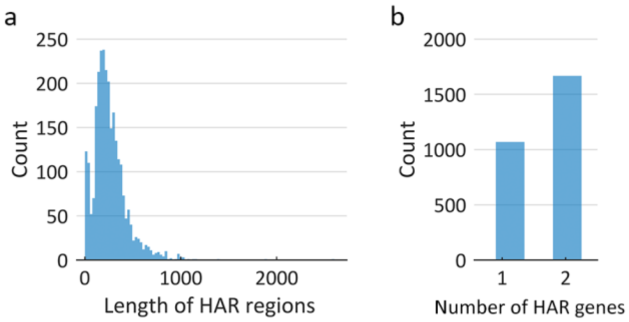


**Supplementary Figure 7.** Distributions of lengths of the HAR regions (a) and the number of genes mapped from each HAR region (b).

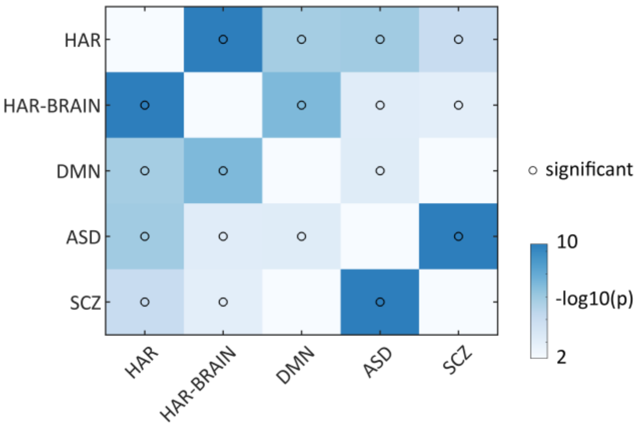


**Supplementary Figure 8.** Enrichment of HAR, HAR-BRAIN, and DMN genes in rare variations of ASD and schizophrenia (SCZ).

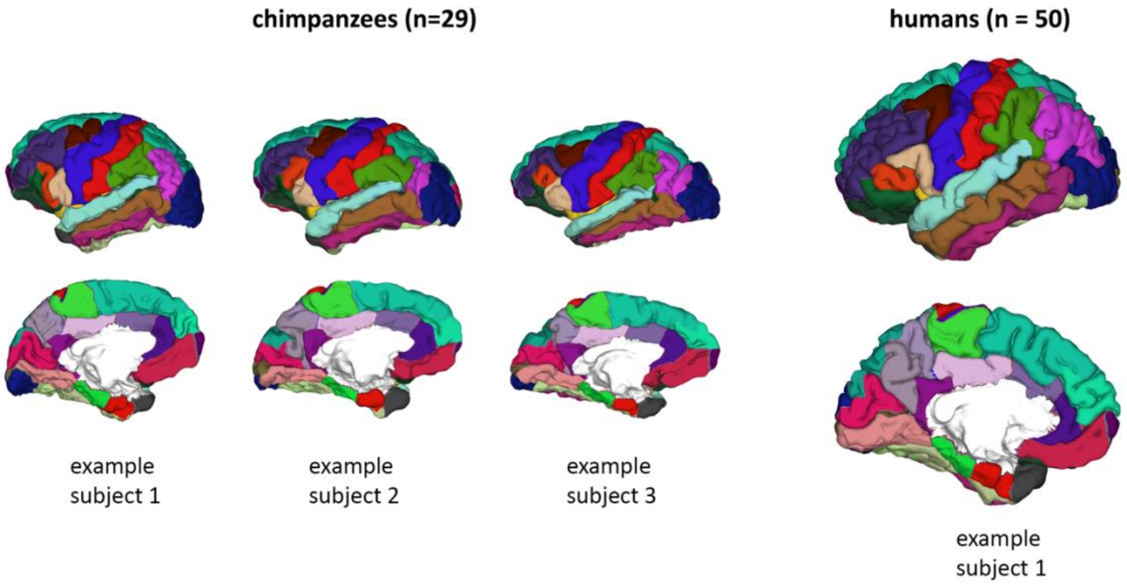


**Supplementary Figure 9.** *Example cortical parcellations of chimpanzees and humans.*

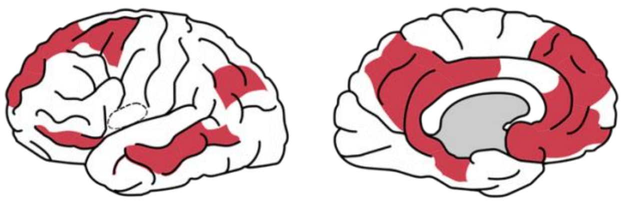

**Supplementary Figure 10.** *DMN assignment adapted from Raichle (2015).*

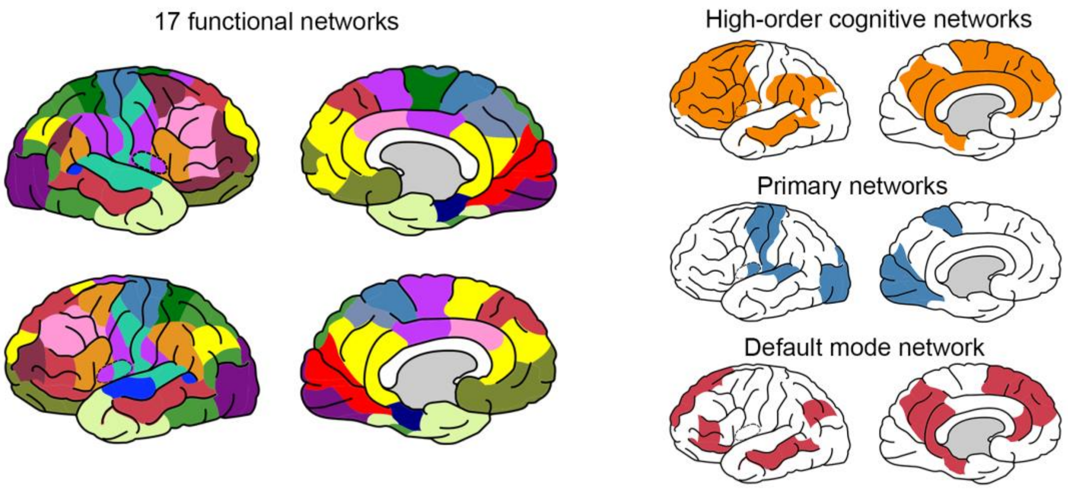


**Supplementary Figure 11.** *Seventeen functional network division.* DK cortical regions were also assigned to 17 functional networks for validation (left panel) [17]. Networks labeled as 11, 15, 16, and 17 were grouped into the default-mode network. Grouping the default-mode network and networks labeled as 7, 8, 12, and 13 formed the higher-order cognitive networks. Networks labeled as 1, 2, 3, and 4 were defined as primary networks (right panel).

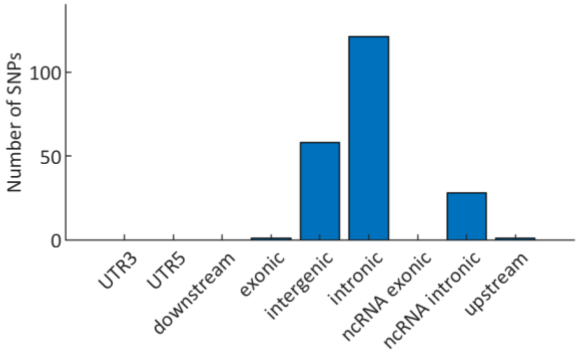


**Supplementary Figure 12**. *Functional annotations of candidate SNPs associated with DMN-FC.*

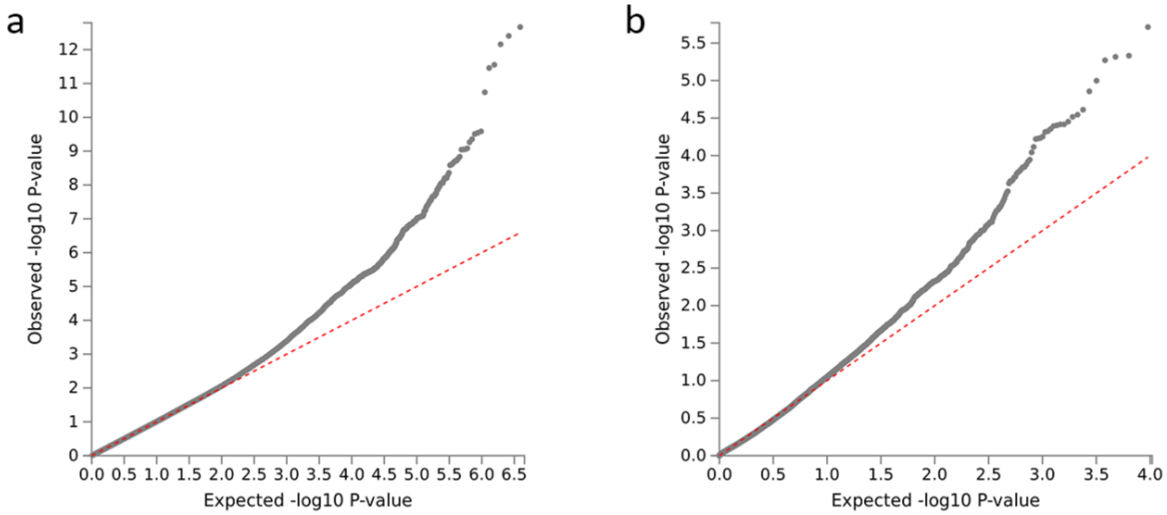


**Supplementary Figure 13**. *QQ-plots of SNP and gene association results in GWAS DMN-FC analysis*. Observed -$log_{10}$ transformed two-tailed *p* values of associations with DMN-FC are plotted against expected null *p* values for a) all SNPs in the GWAS meta-analysis, and b) all genes in the gene-based meta-analysis.
